# Extracellular matrix spatial ecosystems define stem-like niches and predict post-recurrence survival in glioblastomas

**DOI:** 10.64898/2026.11.16.766672

**Authors:** Marina A. Ayad, Ling-Kai Shih, Jiabo Li, Petros Basakis, Hannah Nuszen, Lee A.D Cooper, Daniel J. Brat

## Abstract

Extracellular matrix (ECM) remodeling is a hallmark of glioblastoma (GBM) yet has typically been studied through a reductionist lens, as individual genes or pathways rather than large scale, spatially integrated analyses coordinated with biological states. Focusing on whole matrisome gene expression, we identified three reproducible and spatially coherent ECM transcriptional programs, termed Homeostatic, Stress, and Fibrous, that were associated with specific microenvironmental functions and neoplastic cell states, with the strongest association noted between Homeostatic ECM and neoplastic quiescent stem cell-like (QSC) niches. Ligand-receptor analyses identified FGF1 and BGN as candidate ECM-driven regulators of QSC. In an independent longitudinal bulk transcriptomic cohort, ECM-based models outperformed traditional clinical and biomarker variables for predicting post-recurrence survival, with elevated Homeostatic ECM activity associated with improved post-recurrence survival, while MGMT promoter methylation no longer remained prognostic. Overall, these findings establish ECM organization as a clinically relevant dimension of GBM biology and nominate ECM-associated signaling as candidate therapeutic targets.

## INTRODUCTION

Glioblastoma (GBM) is the highest grade primary brain tumor with approximately 10,000 deaths per year in the United States alone [1]. Although initially therapeutically responsive, recurrence is inevitable, leading to a median overall survival of approximately 16 months and post-recurrence survivals ranging from 6-9 months [2]. Rapid progression is characterized by dramatic shifts in the tumor micro-environment (TME) and dynamic reorganization of the extracellular matrix (ECM) [3]. In healthy brain, ECM provides biochemical and biophysical signals that contribute to neuronal function including synaptic stabilization and neural plasticity, while also maintaining the blood-brain barrier [4, 5]. In contrast, extensive ECM remodeling is a hallmark of glioma progression and described as the “silent architect” [6]. TME restructuring and ECM remodeling are particularly prominent in response to the development of necrosis in GBM, when spatially and biologically distinct zones orient themselves around the necrotic core and promote rapid neoplastic growth outward [7–10]. Immediately adjacent to the necrotic core is a severely hypoxic peri-necrotic zone composed of a high density of neoplastic cells and tumor associated macrophages (TAMs). Next is a zone of high vascular density created by secretion of pro-angiogenic factors by the peri-necrotic region. More peripherally is a diminishing gradient of neoplastic and immune cell density that extends to the infiltrating edge, which dissipates into the native brain. Prior research on neoplastic cell crosstalk with ECM has focused almost exclusively on single gene, single protein, or limited cell-cell interactions that provide only a partial view of the impact of ECM. This reductionist approach falls short in yielding insight into this highly interconnected network and its role in GBM progression and recurrence. More recently, spatially resolved approaches have characterized the heterogeneity and organization of ECM across the GBM TME, highlighting the importance of studying the ECM *in situ,* yet have not fully addressed biological impact [11–13]. We have taken the approach of studying the GBM ECM as a dynamic structure continuously shaped by coordinated contribution of neoplastic, immune, vascular and the native brain parenchyma, with distinct ECM programs associated with key neoplastic hallmarks that include hypoxia, stemness, quiescence, proliferation and adhesion [14, 15].

In this investigation, we used the matrisome database [16] which consists of structural components including collagens, glycoproteins and proteoglycans; matrisome-associated elements consisting of ECM-affiliated proteins such as annexins and ECM regulators, including matrix metallopeptidases; and ECM secreted factors such as interleukins, S100 calcium-binding proteins and fibroblast growth factors [16]. Although the ECM is ultimately composed of proteins deposited spatially over time, transcriptomic profiling adds value by more fully characterizing biological processes that drive ECM organization and niche formation, which is especially relevant in GBM with its rapid evolution, cellular plasticity, and microenvironmental remodeling. We initially focused our factorization on a curated matrisome gene panel to resolve latent organizational states specific to the ECM compartment regardless of the cells expressing them. This ECM-focused decomposition was designed to recover structure within a single biological axis that would otherwise be diluted from genome-wide factorizations dominated by larger sources of transcriptional variance. We next hypothesized that ECM organization itself defines a coherent biological axis of variation not fully explained by cellular composition or histological region. The generation of a data-driven representation of ECM organization moves beyond conventional single enrichment scores that obscures functional heterogeneity and instead identifies recurrent transcriptional programs across tissue while enabling spatial analyses of matrix-tumor interaction.

By applying this holistic framework [7], three spatially coherent ECM transcriptional programs emerged designated as Homeostatic, Stress and Fibrous, with functional TME states including hippo, adhesion, hypoxia, and immune suppression. Within programs we identify biologically relevant neoplastic states, including quiescent stemness (QSC), stemness (GSC), and proliferation, as well as candidate ligand-receptor interactions that regulate them. Finally, using the independent GLASS longitudinal bulk transcriptomic cohort [17], we demonstrate that spatially defined ECM programs have prognostic value post-recurrence that are more predictive than established clinical or molecular features. Collectively, these findings establish a spatial framework linking ECM transcriptional programs, key cancer pathways, tumor cell states, and prognosis (**Fig. 1**).

**Figure 1.**
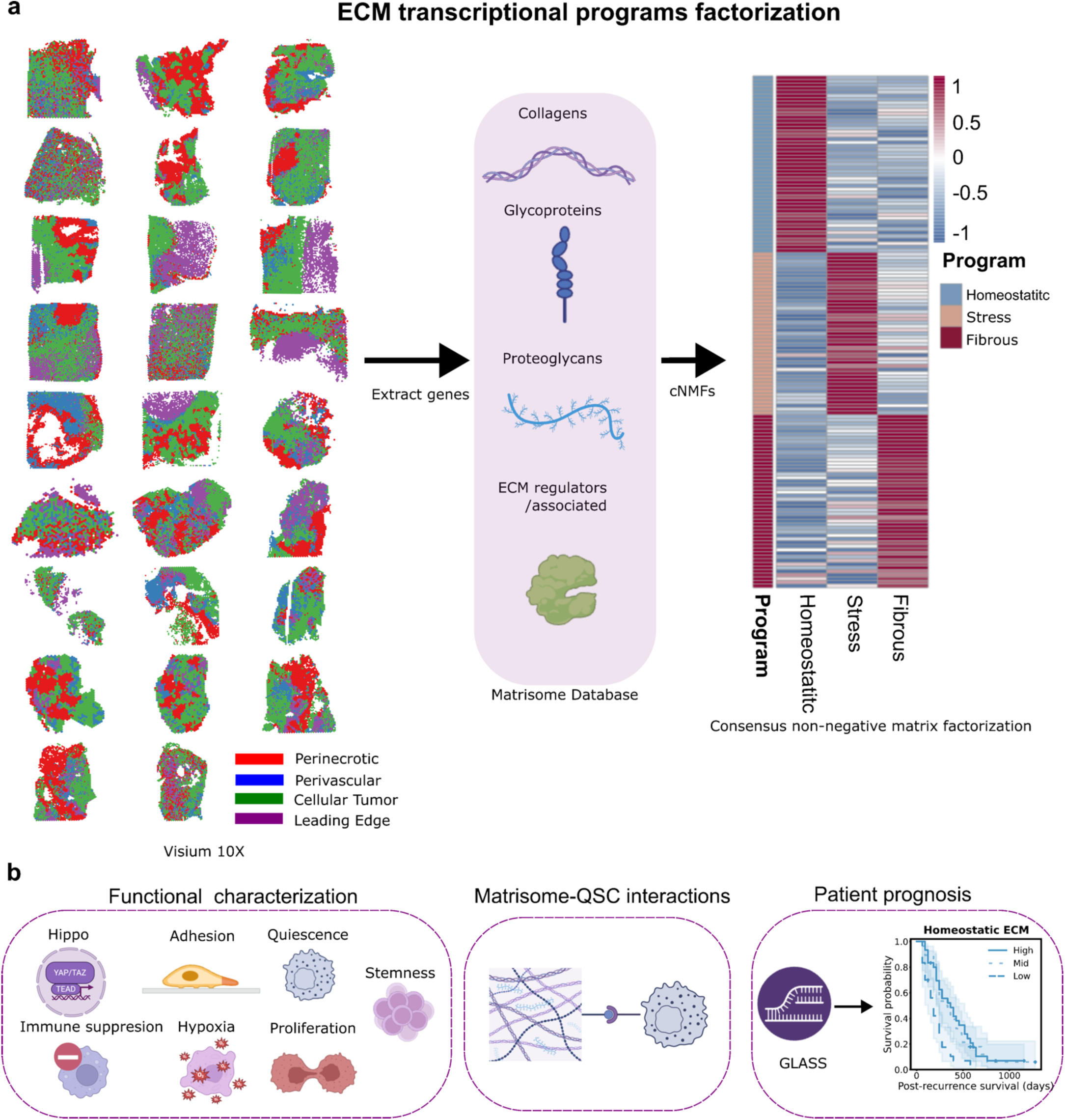
ECM transcriptional program discovery and downstream analysis overview. **(a)** Histological Ivy GAP region annotations across 26 Visium spatial transcriptomics specimen. Matrisome genes are extracted and subjected to consensus non-negative matrix factorization (cNMF), identifying Homeostatic, Stress and Fibrous ECM programs. The heatmap displays the weighted gene contributions defining each transcriptional program. **(b)** Overview of downstream analyses, including functional characterization of ECM programs, ECM-QSC ligand-receptor interaction analysis, and evaluation of ECM program’s prognostic significance on an independent longitudinal bulk transcriptomic cohort. The Kaplan-Meier plot shows post-recurrence survival stratified based on Homeostatic ECM expression levels.

## RESULTS

### cNMF resolves three ECM transcriptional programs in GBM

To identify ECM transcriptional states independent of their cellular sources, we applied consensus non-negative matrix factorization (cNMF) [18] to the curated matrisome gene set. Log-normalized expression data were extracted from all 26 Visium specimens obtained from 18 patients and merged across spots prior factorization (**Fig.1a**). The optimal k was selected using the k-selection diagnostic (**Fig. S1**), balancing reconstruction error against consensus stability. Three ECM transcriptional programs emerged from consensus NMF: Homeostatic, Stress, and Fibrous, each defined by a distinct set of marker genes (**Fig. 2a**). These programs exhibited distinct gene-loading profiles and spot-level activities with limited correlation between programs (**Fig. S1b-c**). The Homeostatic program exhibited strong expression of astrocyte-associated genes, with S100 family (S100B, S100A13, S100A6, S100A1) reflecting calcium signaling. This program was also characterized by proteolytic inhibitors (CST3, and SERPINI1), which have functions consistent with maintenance of tissue homeostasis and relative restraint of extracellular remodeling. The Stress program was marked by angiogenic and tissue-remodeling factors, including VEGFA and ANGPTL4 (hypoxia/angiogenesis), as well as SERPINE1, PLAU, TNC, POSTN, TIMP1 (wound-healing associated matrix remodeling), suggesting an ECM program with an active microenvironmental stress response and remodeling under oxygen deprivation. The Fibrous program was dominated by fibrillar collagens (COL1A1, COL3A1, COL1A2, COL5A1, COL6A2), together with collagen-processing and crosslinking enzymes (LOXL2, PCOLCE, SERPINH1), structural proteoglycans (BGN, HSPG2), and matrix associated genes (FN1, TGFB1, and MMP2). This program was consistent with extensive collagen deposition and matrix organization characteristic of a fibrotic ECM program. These three programs were reproducible across the cohort and were used as the basis for spatial and microenvironmental association analyses.

**Figure 2.**
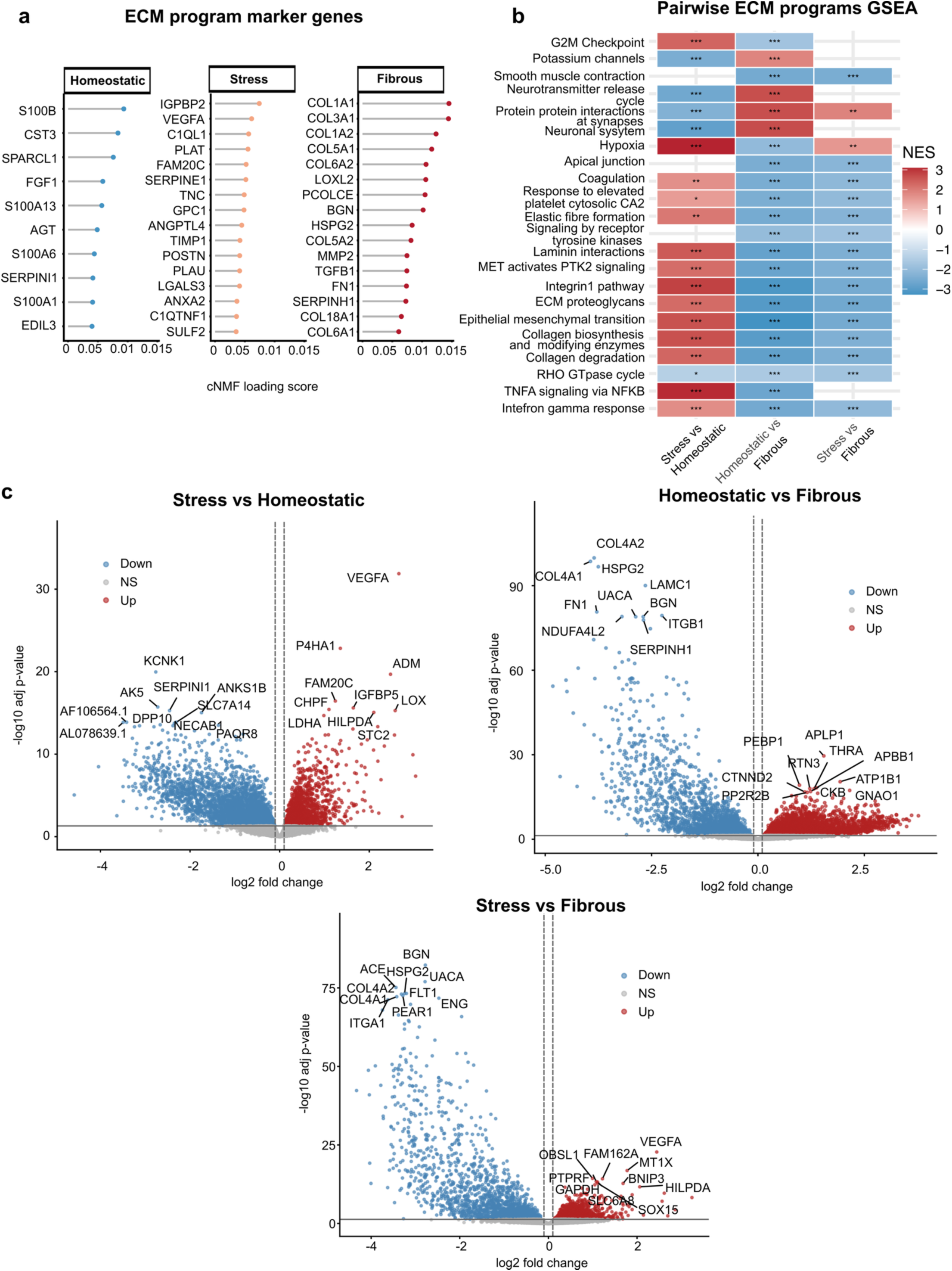
Homeostatic, Stress, and Fibrous ECM programs and associated microenvironmental states. **(a)** Top marker genes defining the Homeostatic, Stress and Fibrous ECM programs. Genes are ranked by their cNMF loading score, which is the weight of each gene in the program’s gene expression signature. **(b)** Pairwise GSEA reveals molecular pathways enriched across ECM programs. Color scale represents normalized enrichment score (NES). Asterisks denote adjusted p-value (* < 0.05, ** < 0.01, *** < 0.001). **(c)** Volcano plots showing pseudo-bulk differential gene expression between ECM programs. Genes up- or down-regulated in the first named program relative to the second are shown in red and blue, respectively; NS, not significant. Dashed lines mark the log2 fold-change cutoffs (±X) and the horizontal line marks adjusted p = 0.05. Representative genes are labeled.

### ECM programs reflect distinct microenvironmental states

We performed transcriptome-wide pseudo bulk differential gene expression between spots dominated by each ECM program. Although the ECM programs were derived from matrisome genes alone, pairwise comparisons revealed transcriptomic differences across the full gene expression landscape (**Fig. 2b**). Differential expression analysis identified 3917 genes between Stress and Fibrous programs, 6453 genes between Homeostatic and Fibrous ECM, and 5610 genes between Stress and Homeostatic ECM (BH corrected, p < 0.05) (**<u>Table S1</u>**) and shown in volcano plots (**Fig. 2c**). The largest transcriptional divergence between programs was observed between Fibrous and Homeostatic ECM, with Stress occupying an intermediate position. These broad transcriptional differences were further investigated using gene set enrichment analysis (GSEA) which was applied across Hallmark, PID, Reactome gene sets comparing Homeostatic, Stress, and Fibrous ECM spots (**<u>Table S2</u>**). Pairwise GSEA revealed five recurring transcriptional themes distinguishing the three ECM transcriptional programs (**Fig 2b**), with extended pathways shown in **Fig. S2**. First, ECM remodeling and matrix organization including collagen formation, degradation, and biosynthesis, ECM proteoglycans, integrin and laminin signaling, MET-PTK2 signaling, epithelial-mesenchymal transition, and the G2M checkpoint, showed a consistent three-way gradient highest in Fibrous, intermediate in Stress, and lowest in Homeostatic ECM (all pairwise BH corrected p < 0.05). Second, Fibrous-dominant spots were additionally characterized with coagulation and vascular-associated signature. Coagulation followed the same pattern of Fibrous, Stress, Homeostatic ECM gradient, while platelet-related signaling pathways were specifically elevated in the Fibrous-dominant spots relative to both other programs without a significant difference between Stress and Homeostatic ECM. Third, hypoxia showed a distinct pattern from the ECM-remodeling and coagulation themes, highest in Stress, intermediate in Fibrous and lowest in Homeostatic ECM. Fourth, inflammatory signaling (TNFA signaling via NF-kB interferon gamma response), was elevated in both Stress and Fibrous relative to Homeostatic ECM. Finally, Homeostatic-dominant ECM spots were enriched for neuronal and synaptic signaling including neurotransmitter release, protein-protein interactions at synapses, and potassium-channel activity, consistent with the preservation of normal brain parenchyma function. Collectively, these findings demonstrate that ECM transcriptional programs capture a coordinated biological axis of microenvironmental variation within the GBM TME.

### ECM transcriptional programs define spatial ecosystems in primary GBM

For each spot, NMF generated a continuous score for all three ECM programs, representing the relative contribution of each program within a spot. ECM programs appeared spatially distinct (**Fig. 3a**) with spatial organization reflected in program-level spatial autocorrelation measured using Global Moran’s I. Across the cohort, all three ECM programs showed moderate to strong autocorrelation (median Moran’s I = 0.66, 0.56, and 0.42 for Homeostatic, Stress and Fibrous ECM respectively; range 0.01-0.87), with 80% of sample-program pairs exceeding Moran’s I > 0.3 (**Fig. 3b**, **<u>Table S3</u>, <u>4</u>**). These findings indicate that spots with similar ECM programs tend to cluster spatially rather than being randomly distributed.

**Figure 3.**
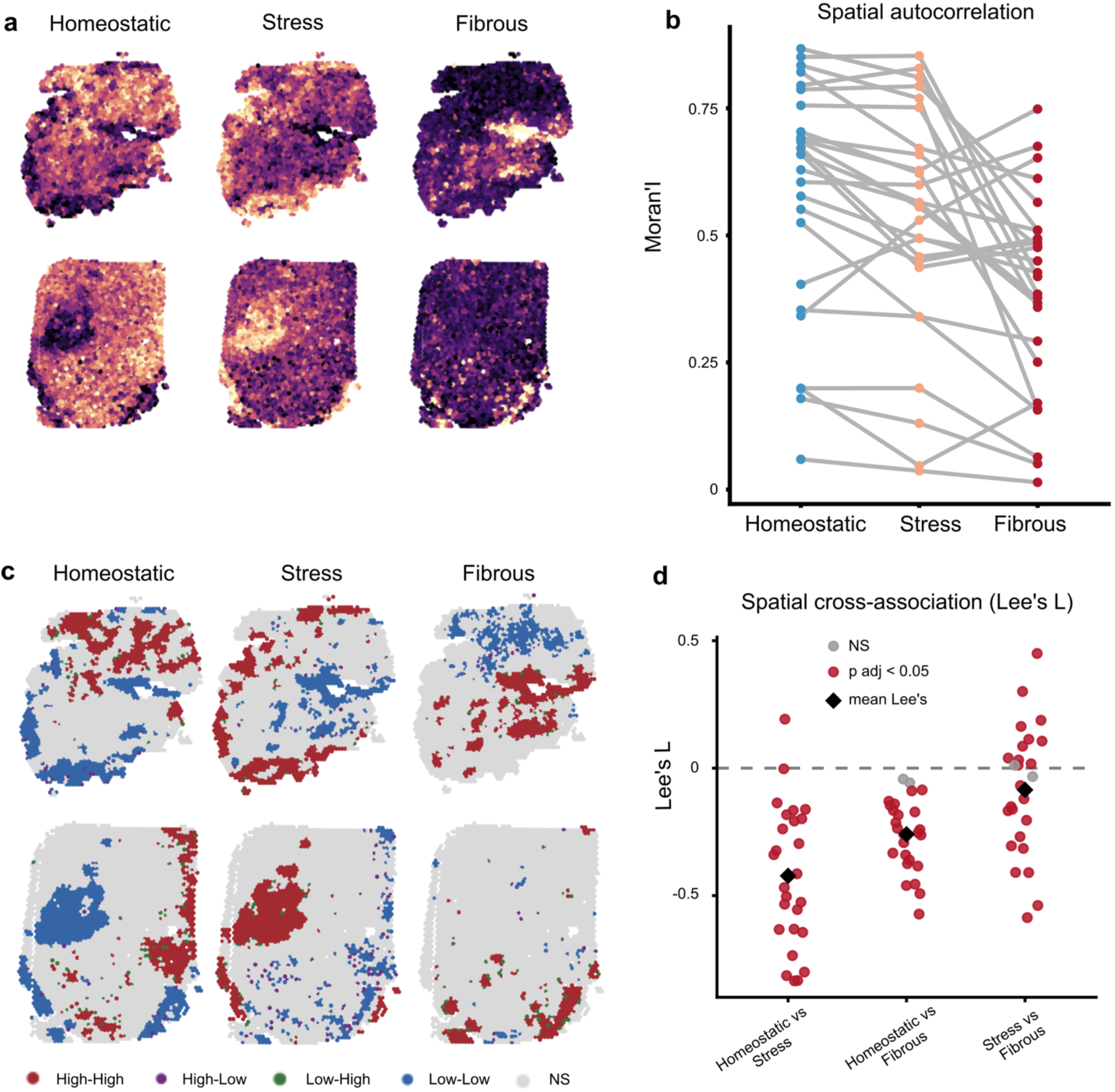
ECM programs exhibit distinct spatial organization. **(a)** Representative spatial distributions of Homeostatic, Stress, and Fibrous ECM program activity in GBM Visium spatial transcriptomic specimens. **(b)** Mean spatial autocorrelation of each ECM program per specimen measured by Moran’s *I*. **(c)** Local spatial association (LISA) maps showing regions of ECM hot spots (red), cold spots (blue), transitioning neighborhoods: from high to low(purple), from low to high(green), non-significant (NS) spatial autocorrelation (gray) **(d)** Pairwise spatial cross-association analysis between ECM programs using Lee’s *L*, demonstrating predominantly negative spatial associations across specimens, indicating spatial segregation. A subset of specimens demonstrated a Stress-Fibrous co-localization. Black diamonds indicate mean Lee’s *L* values.

Local Indicators of Spatial Association (LISA) maps were used to localize where this clustering occurs (**Fig. 3c**). Each spot was classified by comparing its own program activity to that of its neighbors. A high-high (red) spot represents high program activity surrounded by neighbors with similarly high activity, marking the ECM program hotspot. The cold spots (blue) correspond to low ECM program activity surrounded by similarly low activity. The scarcity of the high-low (purple) and low-high (green) indicates limited sharp transitions between ECM transcriptional programs. Both Homeostatic and Stress ECM formed larger hotspots across specimens, while the Fibrous program formed more spatially confined hotspots (**Fig. S3**).

Lee’s L statistic was used to calculate the spatial association between pairwise ECM program scores capturing both the strength and directionality of co-patterning. A negative Lee’s L indicates a spatial segregation between the paired programs. Homeostatic vs Stress and Homeostatic vs Fibrous ECM showed a consistent and strong segregation across specimens (median Lee’s L = -0.44, and -0.25, respectively), while Stress vs Fibrous did not consistently segregate between specimens (median Lee’s L = -0.05) (**Fig. 3d**).

The variation in Lee’s L directionality between Stress and Fibrous programs was not random, rather, it was strongly anti-correlated with the degree of Homeostatic-Stress ECM segregation across the same specimens (Pearson r = -0.72, p-adj = 8.5e-5). This association was specific to this pair combination where neither Homeostatic-Fibrous ECM vs Stress-Fibrous ECM (r=-0.18, p-adj=0.55), nor Homeostatic-Fibrous ECM (r=0.09, p-adj=0.65) showed a comparable association (**<u>Table S5</u>**). This pattern is reflected in the two main clusters of the dendrogram (**Fig. S4**). Specimens with the strongest Homeostatic-Stress ECM segregation (negative Lee’s L) showed Stress-Fibrous ECM co-localization, while specimens with weak Homeostatic-Stress ECM segregation showed higher segregation between Stress-Fibrous. These findings suggest that ECM programs are arranged into coordinated spatial ecosystems rather than acting as independent transcriptional states.

### ECM transcriptional programs capture hallmark GBM microenvironmental niches

ECM transcriptional programs were associated with key functional pathways in GBM, including hypoxia, immune suppression, Hippo signaling, and adhesion. A univariate linear mixed model fitted to predict these signatures from ECM transcriptional programs revealed the strongest positive association between Fibrous ECM and adhesion (β = 0.61, R^2^m = 0.27), and between Stress ECM and Hypoxia (β = 0.57, R^2^m = 0.26). Glioma Stem Cell association was observed between the Homeostatic ECM and Hypoxia signature (β = -0.63, R^2^m = 0.31). Fibrous ECM was also strongly associated with the Hippo signature (β = 0.44, R^2^m = 0.15) (**Fig. 4a**, **<u>Table</u> <u>S6</u>**)

**Figure 4.**
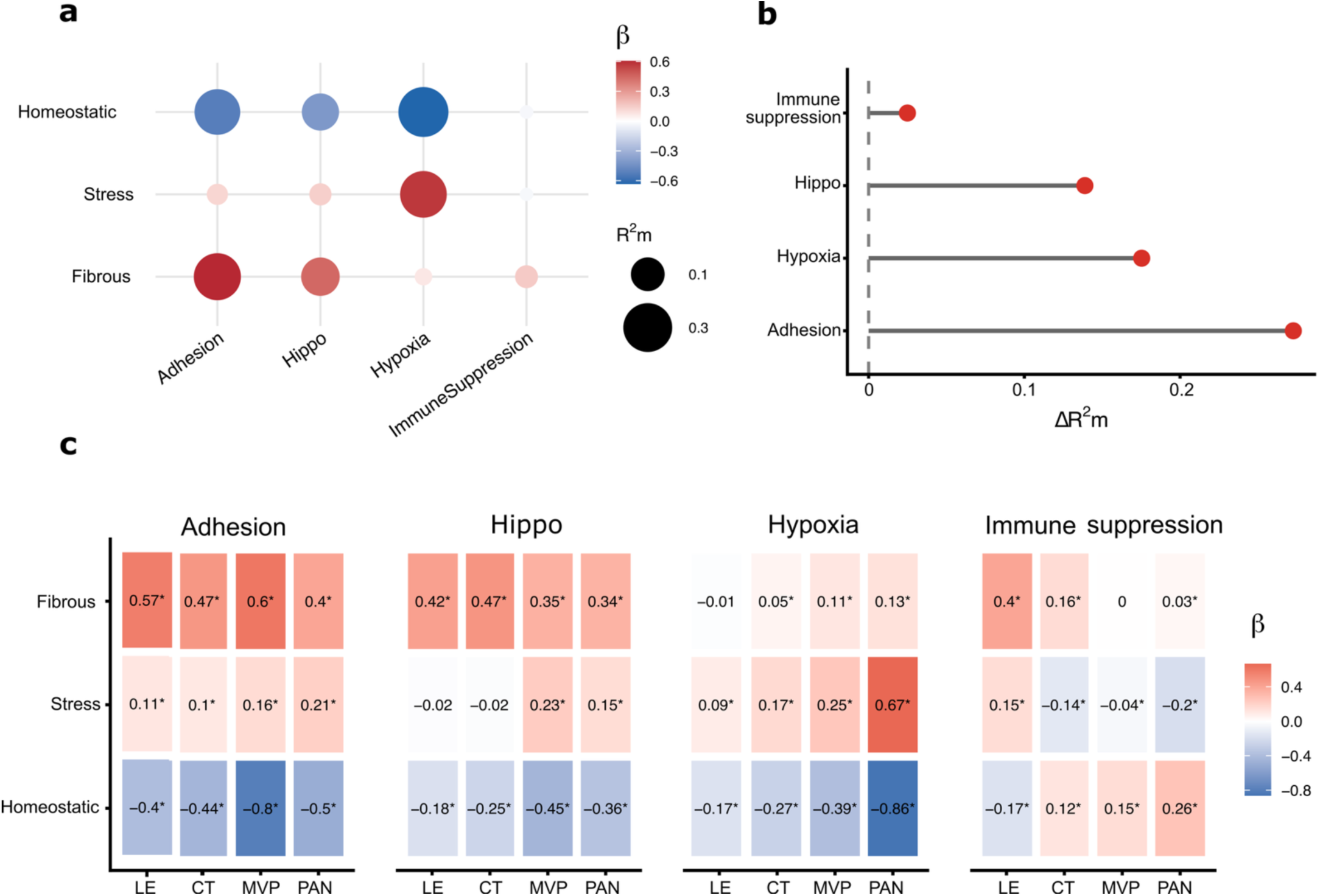
Associations between ECM programs and GBM hallmark niches. **(a)** Associations between ECM program activity and hallmark niche signatures (adhesion, Hippo, hypoxia, and immune suppression) estimated using linear mixed-effects model. Circle color indicates the regression coefficient (β), and circle size represents the marginal variance explained (R²m). **(b)** Incremental variance explained (ΔR²m) by ECM programs beyond histological regions for predicting Adhesion, Hippo, Hypoxia, and Immune suppression. **(c)** ECM-hallmark niche interactions across histological regions. Heatmap colors indicate regression coefficients (β) describing the association between ECM program activity and ECM-cell state interactions with each histological region. Asterisks denote statistically significant associations (p < 0.05). Histological regions defined by Ivy GAP include Cellular Tumor (CT), Leading Edge (LE), Microvascular Proliferation (MVP), and Pseuodpalisading and Peri-necrotic (PAN) region.

ECM programs explained additional variation beyond conventional Ivy GAP histological classification (Pseudopalisading and Peri-necrotic [PAN], Microvascular Proliferation [MVP], Cellular Tumor [CT], Leading Edge [LE]). Across all four signatures, models incorporating ECM programs provided a better fit than histology-only models, as indicated by lower Bayesian Information Criterion (BIC) values across all four signatures (ΔBIC = 825 - 6899, **<u>Table S7</u>**), with the largest improvements for adhesion (ΔBIC = 6899) and hypoxia (ΔBIC = 6,057). Gains in variance explanation were observed for Adhesion, Hypoxia, Hippo, and Immune Suppression (ΔR^2^m = 0.272, 0.175, 0.139, 0.025, **Fig. 4b, Fig. S5**).

To determine whether ECM program-signature associations vary by histological region, we tested for an ECM × Ivy GAP region interaction for each signature, accounting for the nested patient and specimen structure. All twelve interactions were significant (likelihood ratio test, BH adjusted, all p < 0.0001, **<u>Table S8</u>**, see Methods), with strongest evidence of histology-dependent ECM associations for the hypoxia signature (**Fig. 4c**). Homeostatic ECM showed a negative association with Adhesion, Hippo and Hypoxia across all regions, strongest in PAN for Hypoxia (β = -0.86, and in perivascular regions for Adhesion and Hippo (β = -0.8, -0.45, respectively) while showing a positive association with Immune Suppression most pronounced in the PAN (β = 0.26). Furthermore, Fibrous ECM showed a consistent positive association with Adhesion and Hippo signaling across all four regions (β = 0.34-0.57). Similar to the Homeostatic in PAN region, Fibrous ECM activity was low in the leading edge overall, and this steep slope may partly reflect a restricted dynamic range rather than a uniformly strong coupling (**<u>Table S9</u>**). Stress ECM showed the most region dependent behavior, its association with Hypoxia was weak in cellular tumor and leading edge (β = 0.17, 0.09) but substantially stronger in PAN (0.67), consistent with the known localization of hypoxic stress within necrotic regions. These results are also reflected in the trends visualized between ECM program tertile usage (Low, Medium, High) and every signature (**Fig. S6**). Overall, these results reflect that ECM transcriptional programs provide biological information beyond conventional histological classification. Nevertheless, the strength of these relationships remained context-dependent and was most pronounced from the association between Stress ECM and hypoxia within the PAN region.

### ECM transcriptional programs are associated with distinct neoplastic cell states

To identify associations between ECM transcriptional programs and neoplastic cell states, the analysis was restricted to neoplastic spots. Quiescent stem cell (QSC), Glioma Stem Cell (GSC) and proliferation signatures were quantified in neoplastic spots (see **Methods**).

A univariate linear mixed model fitted to predict these signatures from ECM transcriptional programs revealed the strongest positive association between Homeostatic ECM expressing spots and QSC (β = 0.43, R^2^m = 0.2) and negative association with Stress ECM (β = -0.35, R^2^m = 0.14) (**Fig. 5a, <u>Table S10</u>**). Similarly, GSC was positively associated with Homeostatic ECM (β = 0.32, R^2^m = 0.09) and negatively with Stress ECM (β = -0.24, R^2^m = 0.055). As for proliferation the associations were weak across ECM programs (R^2^m = 0.001-0.044).

**Figure 5.**
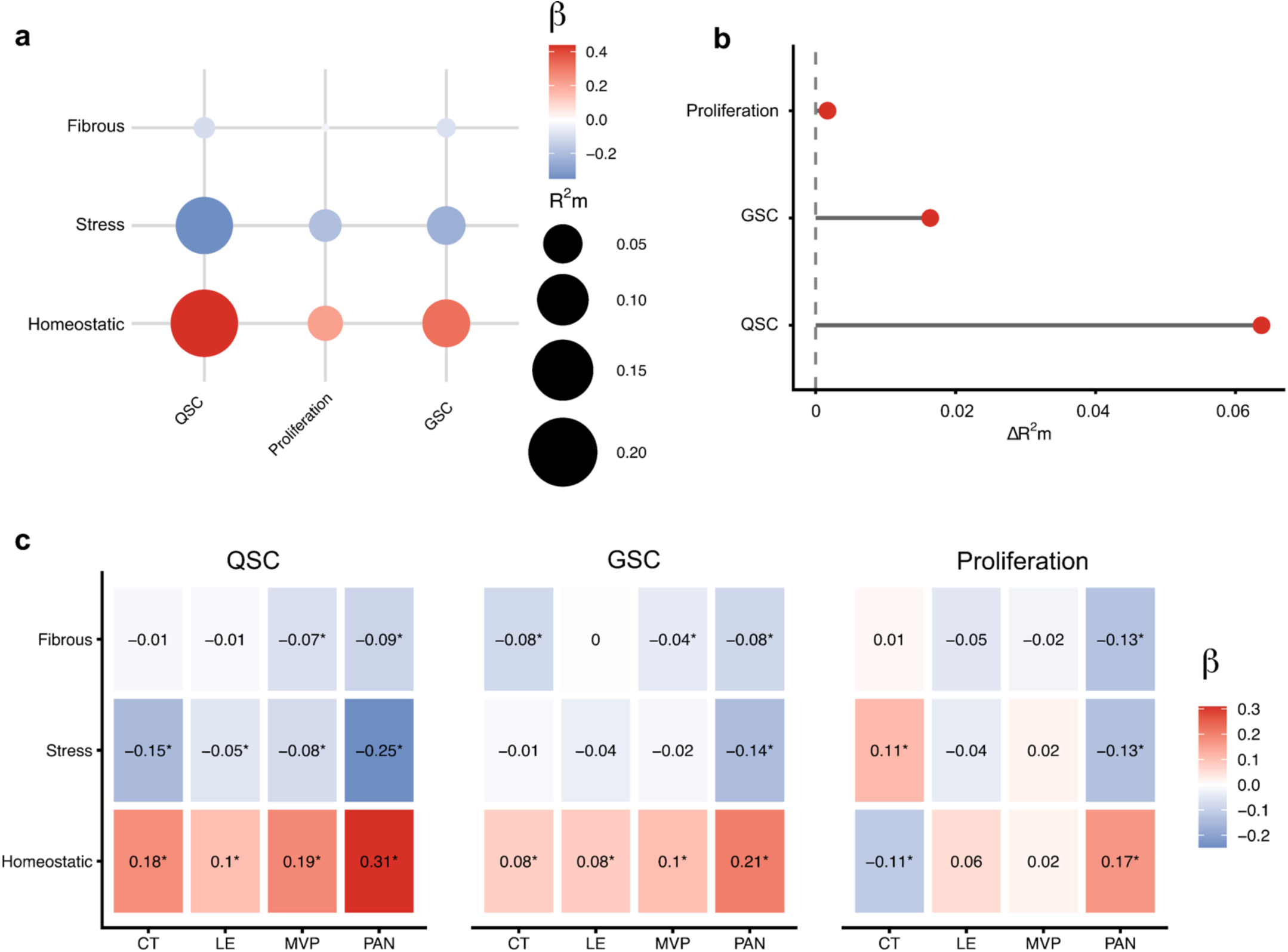
Associations between ECM programs and neoplastic cell states. **(a)** Associations between ECM program activity and neoplastic cell states (QSC, proliferation, GSC) estimated using linear mixed-effects model. Circle color indicates the regression coefficient (β), and circle size represents the marginal variance explained (R²m). **(b)** Incremental variance explained (ΔR²m) by ECM programs beyond histological regions for predicting QSC, GSC, and proliferative cell states. **(c)** ECM-cell state interactions across histological regions. Heatmap colors indicate regression coefficients (β) describing the association between ECM program activity and neoplastic cell states within each histological region. Asterisks denote statistically significant associations (p < 0.05).

To determine whether ECM programs capture neoplastic cell states beyond conventional histological classification, we compared mixed-effects models containing Ivy GAP regions alone or in combination with ECM programs. Across all three signatures, models incorporating ECM programs provided a better fit than histological-only models, as indicated by BIC (**Fig. 5b, <u>Table S11</u>, Fig. S7**). The largest improvements were observed for QSC (ΔR^2^m = 0.064, ΔBIC = 575.68) and GSC (ΔR^2^m = 0.016, ΔBIC = 94.53). However, only a minimal improvement was observed for proliferation (ΔR^2^m = 0.002, ΔBIC = 10.68). This implies that ECM programs capture more variation in stem-like states than in proliferative activity.

Consistent with the expected inverse relationship between quiescence and proliferation, QSC and proliferation scores were negatively correlated in the CT, LE, and MVP regions (Pearson’s r = -0.22, -0.23, -0.24, respectively) (**Fig. S8, <u>Table S12</u>**). This anticorrelation was absent in the PAN region (Pearson’s r = -0.008), suggesting that the relationship between quiescence and proliferation varies across histological regions and may be influenced by region-specific ECM programs.

To determine whether ECM program-signature associations vary by histological region, we tested for an ECM × Ivy GAP region interaction for each signature, accounting for specimens as a random effect. All 9 interactions were significant (BH-adjusted likelihood ratio test, p = 1.5×10^-44^ - 0.04, **<u>Table S13</u>**). The interaction was strongest for proliferation with both Homeostatic and Stress ECM programs, driven by a sign reversal across histologic regions. Homeostatic ECM activity was associated with lower proliferation within cellular neoplastic spots (β = -0.11, p <0.05), but with higher proliferation within the PAN region spots (β = 0.17, p <0.05). Stress ECM showed the opposite pattern, associated with higher proliferation in cellular tumor spots (β = 0.11, p <0.05) but lower proliferation in the PAN zone (β = -0.13, p <0.05) (**Fig. 5c**). By contrast, Homeostatic ECM was consistently positively associated with higher quiescence across all four regions, with the association strengthening toward the PAN zone (β = 0.1-0.31, all p < 0.05), and showed a similarly consistent but weaker positive association with stemness across all regions (β = 0.08 - 0.21, p< 0.05). These trends were consistent with these results when ECM programs were instead calculated as discrete tertiles (Low, Medium, High) rather than continuous scores (**Fig. S9**). Overall, the strongest association was observed between the Homeostatic program and QSC state, with ECM programs explaining additional variation beyond that captured by histological region classification.

### ECM programs identify FGF1 and BGN as candidate regulators of QSC state

Since QSC state showed the strongest associations with ECM programs, NicheNet [19] was used to identify upstream ECM-derived ligands predicted to regulate QSC states. NicheNet analyses were performed independently for each ECM program using ECM-high spots as senders and QSC-high neoplastic spots as receivers. NicheNet ligand activity was then evaluated using a consensus QSC gene set derived from differential expression between high and low tertiles of QSC neoplastic spots (**Fig. S10**). Candidate ligands ranked by predicted activity included FGF1, EDIL3, and S100B from Homeostatic ECM spots. TIMP1, ANGPTL4, TNC, and VEGFA from Stress ECM spots, and COL3A1, and TGFB1 from Fibrous ECM spots (**Fig. S11, <u>Table S14</u>**).

To identify ligands that are both spatially plausible and consistently enriched across local neighborhoods, a neighborhood-radius sweep was performed of predicted ligand-receptor interactions (**Fig. S12, <u>Table S15</u>, <u>16</u>**). This filtering step prioritized FGF1 from the Homeostatic ECM, and BGN from the Fibrous ECM as the most robust candidate ligands (**Fig. 6a, <u>Table S17</u>**). Predicted interactions included FGF1 signaling through FGFR1, FGFR2, FGFR3, and BGN signaling through TLR2 and TLR4 expressed by in QSC neoplastic spots (**Fig. S13**).

**Figure 6.**
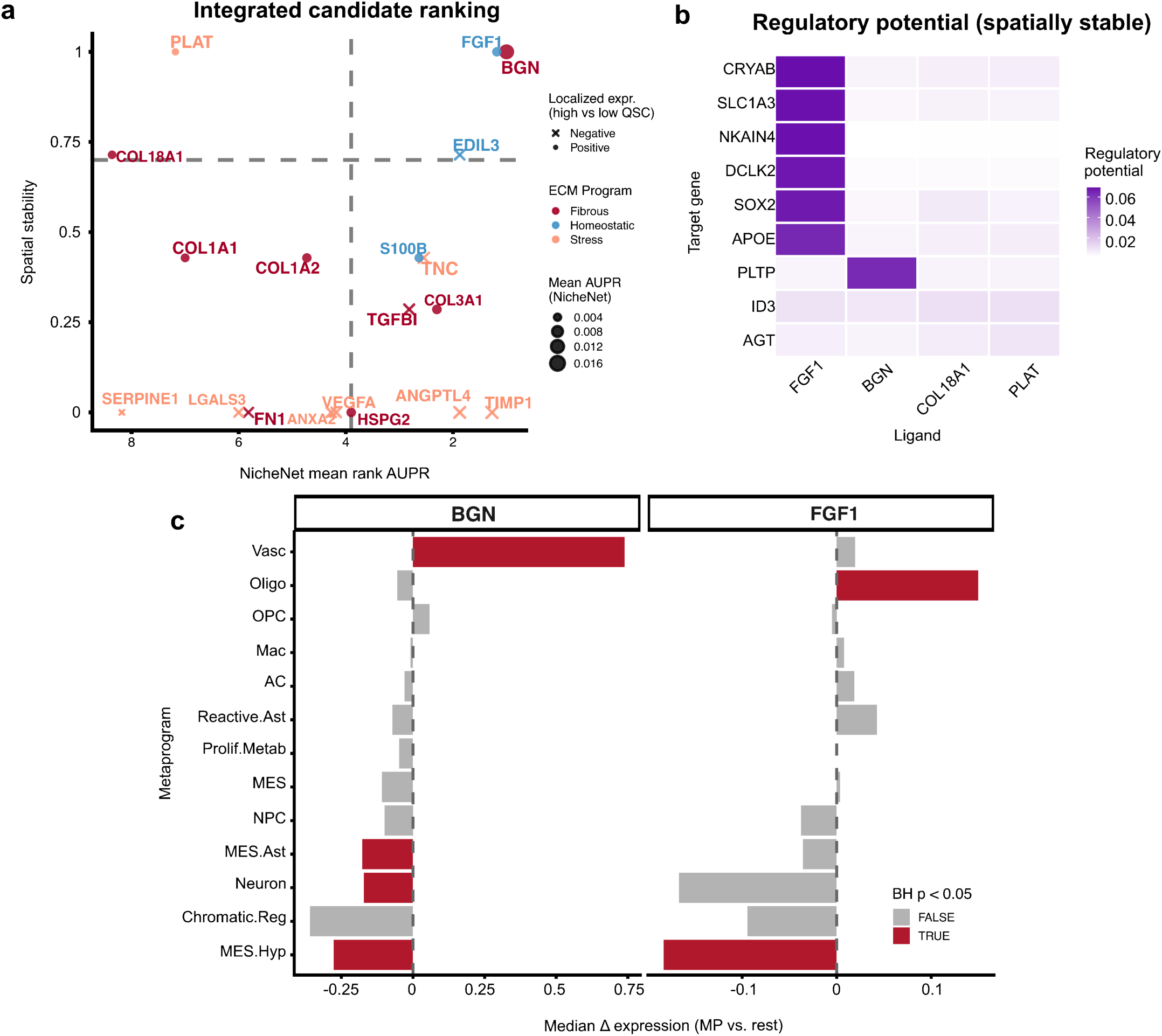
Identification of FGF1 and BGN as candidate mediators of ECM-QSC interactions. **(a)** Integrated ranking of candidate ECM-QSC ligand-target interactions prioritized based on NicheNet regulatory activity, spatial stability across specimens, and differential expression between high- and low QSC neighborhood spots, with high per ECM program as sender and high QSC as receiver spots. Point size indicates mean NicheNet AUPR, color indicates ECM program sources, and symbols indicate the direction if ligand expression in high vs low QSC regions. **(b)** Regulatory potential of prioritized ligands for spatially stable target genes. Heat map colors indicate NicheNet regulatory potential scores between candidate ligands and downstream target genes. (c) Median difference in expression of Metaprogram expression of FGF1 and BGN versus rest of metaprograms reflecting metaprograms FGF1 and BGN-expressing. Red bars denote significant enrichment (BH corrected p<0.05)

NicheNet target predictions suggested that FGF1 may regulate CRYAB, and BGN may regulate PLTP in QSC high spots (**Fig. 6b**). To contextualize the cellular sources associated with these ligands, expression of the transcriptional metaprogram defined by Greenwald et al. was quantified [7] within ligand-associated ECM programs. Across specimens, the oligodendrocyte metaprogram was significantly associated with FGF1 (BH-corrected p= 0.0015), and the vascular program was significantly associated with BGN (p= 4.65×10^-6^), based on one-sample Wilcoxon signed-rank test with BH correction for multiple comparisons (**Fig. 6c, <u>Table S18</u>**). These findings identify FGF1 and BGN as spatially supported ECM-derived signaling candidates linked to neoplastic QSC regions.

### ECM transcriptional programs demonstrate prognostic relevance in GBM

To assess clinical relevance, ECM programs were computed in the GLASS cohort (n=59, matched primary, TP-recurrent pairs, n=58, recurrence R1-overall survival OS, **<u>Table S19</u>**). For time to recurrence, prognostic performance was primarily driven by MGMT promoter methylation status, which was independently associated with a shorter time to recurrence in both the clinical model (HR = 3.92, p= 1.6 × 10^-4^) and combined clinical and ECM models (HR= 4.1, p= 2.3 × 10^-5^) (**Fig. S14a**). ECM programs provided a modest additional prognostic information beyond clinical variables increasing the cross-validated C-index from 0.62 ± 0.06 to 0.63 ± 0.05. Longitudinal analysis of matched primary-recurrent pairs demonstrated enrichment of Homeostatic ECM at recurrence (median Δ = 0.059, FDR-adjusted *p* = 0.008, **Fig. S14b**), while Stress and Fibrous ECM did not show significant temporal changes. The enrichment of Homeostatic ECM suggests a potential treatment-associated shift in ECM state composition.

Moreover, the prognostic relevance of ECM programs persisted following recurrence and became more pronounced in the recurrent setting. MGMT methylation was no longer associated with post-recurrence survival (HR=1.2, p=0.53), while Homeostatic ECM remained protective (HR= 0.65, p= 0.004, **Fig. 7a).** ECM-only models outperformed clinical variables in predicting post-recurrence survival in five-fold cross validation (C-index: ECM-only 0.67 ± 0.11 vs. Clinical-only 0.47 ± 0.17, **Fig. 7b, <u>Table S20</u>**). Kaplan-Meier analysis of post-recurrence survival showed significant stratification by Homeostatic ECM program activity, with patients in the lowest tertile demonstrating poorer survival (log-rank p= 4.5 × 10^-4^, **Fig. 7c**). Stress ECM activity showed a trend toward worse outcomes, but neither Stress nor Fibrous ECM programs were significantly associated with post-recurrence survival (p= 0.18 and p= 0.34 respectively, **Fig. S14c, <u>Table S21</u>**).

**Figure 7.**
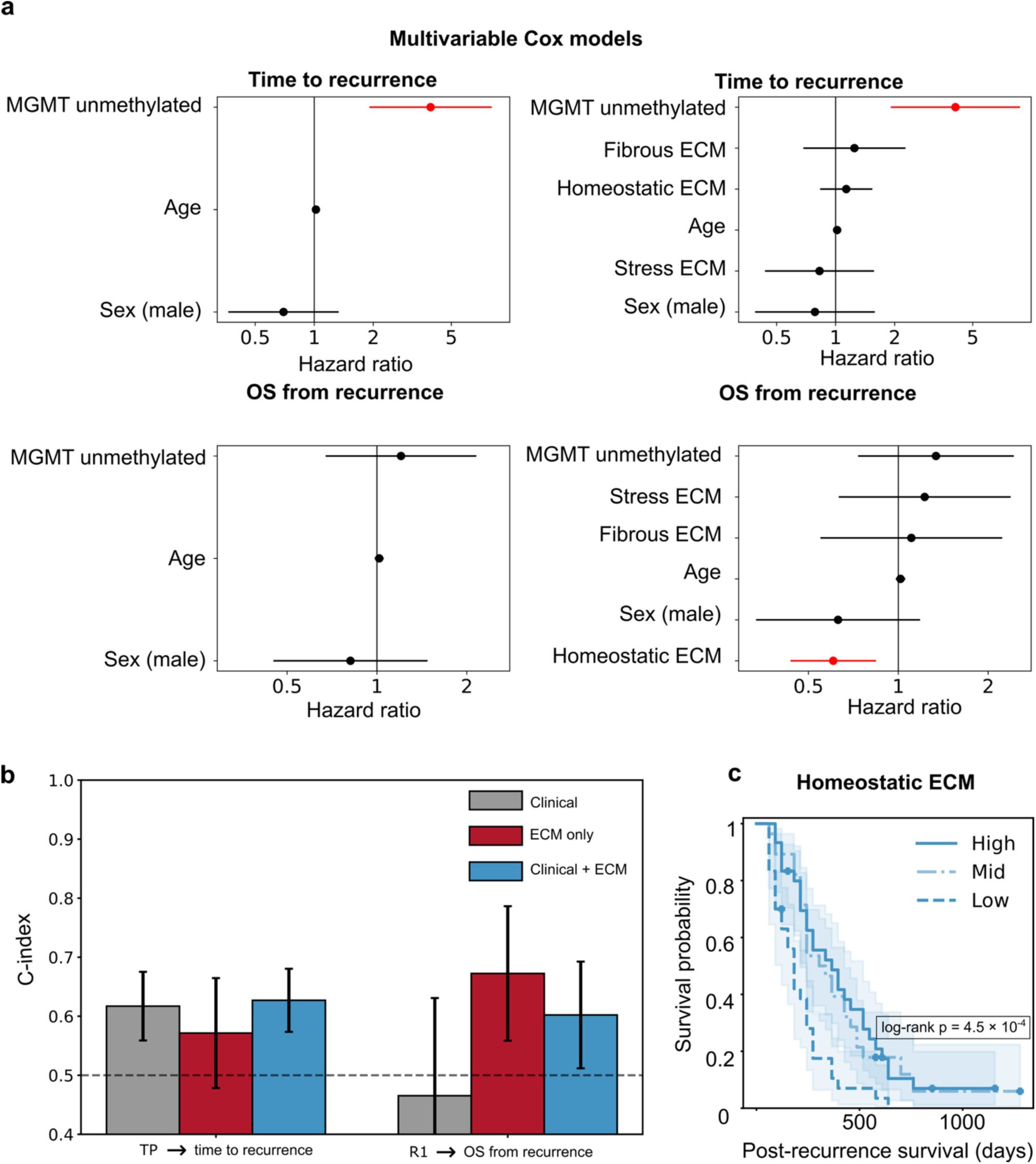
Homeostatic ECM predicts improved post-recurrence survival in recurrent GBM. **(a)** Multivariable Cox proportional hazards models for time to recurrence and post-recurrence overall survival. Models included clinical covariates (age, sex, and MGMT methylation status) with or without ECM programs. Points indicate hazard ratio (HR), and bars denote 95% confidence intervals. Red line denotes statically significant association (P <0.05). **(b)** Mean predictive performance of five-fold clinical, ECM-only and combined models for time to recurrence and post-recurrence survival, measured by concordance index (c-index), bars indicate C-index ranges across the five-fold. **(c)** Kaplan-Meier plot of post-recurrence survival stratified by tertiles od Homeostatic ECM expression. Shaded regions indicate 95% confidence intervals.

## DISCUSSION

Glioblastoma remains one of the most aggressive primary brain tumors and recurrence following initial treatment is nearly universal, reflecting the persistence of therapy-resistant neoplastic cell populations that trigger regrowth. Understanding biologically actionable pathways and mechanisms that sustain residual and recurrent cell populations is essential for developing more effective therapies. Spatial transcriptomic studies have been highly impactful for understanding key axes of the GBM TME, including angiogenesis, invasion and immune cell organization. While ECM remodeling is a hallmark of cancer, its spatial organization and contribution to glioma biology are less appreciated, with only limited studies on individual genes, pathways, or cell types rather than as a coordinated ecosystem. Our findings suggest that the ECM is not merely a passive structural scaffold but rather a biologically active component of the TME organized into distinct transcriptional states [20]. Using cNMF analysis applied to matrisome genes, including core components (collagens, proteoglycans, glycoproteins), matrisome-associated, and secreted factors across 26 specimens from 18 patients, we identified 3 ECM transcriptional programs with distinct spatial organization. These programs represent an additional axis of biological organization within GBM and provide a basis for investigating how ECM programs contribute to tumor progression, therapeutic resistance, and patient outcomes.

The three ECM transcriptional programs were classified by their key genes as Homeostatic, Stress, and Fibrous. The Homeostatic program was enriched for genes associated with normal brain parenchyma with limited remodeling. In contrast, the Stress program was characterized by genes linked to hypoxia and tissue remodeling (VEGFA, and TNC), consistent with a reactive microenvironment. The Fibrous program was enriched for fibrillar matrix components, including several collagens and matrix-crosslinking factors such as LOXL2 and BGN, likely indicating the formation of a dense ECM that may promote growth factor sequestration.

ECM transcriptional programs were spatially organized rather than randomly distributed within the GBM TME, corresponding to distinct environmental states including tissue maintenance, fibrosis, and active remodeling. Notably, Homeostatic and Stress programs consistently occupied segregated zones as reflected by Lee’s L analysis, suggesting that GBM regions are not characterized by a single continuum of ECM remodeling but instead contain competing ecosystems. In contrast, the Fibrous program occupied a more variable spatial relationship colocalizing with Stress in some specimens while remaining segregated with others. Interestingly, this variability was linked to the degree of Homeostatic-Stress segregation. Tumors with strong Homeostatic-Stress segregation had higher Fibrous-Stress colocalization, while tumors with weaker Homeostatic-Stress segregation had greater Fibrous-Stress segregation. This pattern suggests that tumors organize their ECM spatial architecture along one of two competing axes rather than segregating all three programs simultaneously. One spatial mode shows a Homeostatic ECM that is walled off from the rest of the tissue, while Stress and Fibrous remain spatially intermixed, and another mode the Homeostatic program forms a less distinct boundary allowing higher segregation between the Stress and Fibrous spatial domains. One possibility is that fibrotic matrix remodeling reinforces stress-associated compartmentalization.

Beyond defining spatially distinct ECM ecosystems, ECM transcriptional programs were associated with a robust readout of GBM functional states and explained variation in adhesion, hypoxia, Hippo signaling, and immune suppression beyond that reflected by histological localization. While the Fibrous ECM showed the strongest associations with adhesion and Hippo signaling, the relationship between the Stress program and Hypoxia was also strongly tied to the pseudopalisading and peri-necrotic regions, indicating that some ECM-associated functions remain strongly dependent on the local tissue architecture.

More notably, ECM programs were strongly associated with neoplastic cell states. Homeostatic ECM demonstrated the strongest and most consistent relationship with QSC, while Stress ECM showed the opposite pattern, suggesting that QSC preferentially localize within ECM environments resembling tissue maintenance rather than active remodeling. Importantly, the QSC signature used in this study was from an F3-derived transcriptional program previously shown to identify experimentally validated QSCs that remain largely non-proliferative but have the capacity to re-enter the cell cycle following therapeutic stress. This observation links the Homeostatic ECM to a cellular state with established tumor persistence and possible recurrence.

Consistent with prior studies, the QSC signal was anti-correlated with G2-M proliferation across most histological regions. Surprisingly this relationship was lost within the peri-necrotic region, where QSC and proliferation became uncoupled. This finding suggests that the peri-necrotic microenvironment may alter the conventional relationship between QSC and proliferation, potentially reflecting increased cell plasticity. A transitional quiescent-proliferative population has been reported after TMZ treatment [21]. Our data raises the possibility that spatial context may similarly influence the coupling between these two transcriptional states in untreated GBM.

Our ligand-receptor analyses identified FGF1 from the Homeostatic ECM and BGN from the Fibrous ECM as the spatially robust ECM-derived signaling candidates associated with QSC-enriched neoplastic regions. Furthermore, FGF1-associated niches were enriched for oligodendrocyte transcriptional metaprograms, while BGN-associated niches were enriched for vascular metaprograms. It is important to note that NicheNet has limitations when applied to the matrisome, as many structural ECM transcripts, including collagens, are not well represented within its trained regulatory networks. Consequently, these analyses prioritize secreted factors and the matrisome associated transcripts. Overall, the observed association between FGF1-associated niches and oligodendrocyte metaprogram niche suggests that oligodendrocyte-lineage cells may contribute to the maintenance of QSC-associated microenvironments. This interpretation is supported by a prior study demonstrating that FGFR signaling is a critical regulator of glioma stem cell fitness and therapeutic resistance [22]. Moreover, pharmacological inhibition of FGFR signaling has been shown to sensitize GSC to Tumor Treating Fields (TTFields), further highlighting the importance of this pathway in stem-like glioma states [22]. Given that quiescence is a recognized mechanism of therapeutic resistance in glioma, the association of QSCs with predicted FGF1-FGFR signaling raise the possibility that this pathway contributes to maintenance of a QSC state. Under this model, inhibition of FGFR signaling could sensitize stem-like glioma cells to TTFields through disruption of the quiescence-associated programs.

Finally, we examined the prognostic relevance of ECM transcriptional programs in GBM. Although prognostic markers in GBM remain limited due to uniformly poor survival, MGMT promoter methylation status is among the most established [23, 24]. MGMT encodes a DNA repair enzyme that counteracts the effects of TMZ, consequently, tumors with unmethylated MGMT promoters generally exhibit a greater capacity for DNA repair, thereby reducing sensitivity to TMZ [25]. Using Cox survival models incorporating clinical variables including age, sex, and MGMT status, we compared the prognostic value of ECM transcriptional programs quantified from GLASS bulk transcriptomic data which revealed ECM programs had higher prognostic value following recurrence, while MGMT lost its prognostic effect at that disease stage. Stress ECM cases trended toward poorer overall survival, while higher Homeostatic ECM was associated with significantly improved survival outcomes. These findings suggest that ECM programs may not only serve as a prognostic marker but also as therapeutic targets in GBM.

An apparent paradox that emerges from our findings is that although the Homeostatic ECM program was mostly associated with QSCs, elevated Homeostatic ECM expression was also associated with improved survival following recurrence. This apparent paradox may be resolved by distinguishing treatment resistance from prognosis. Quiescence implies resistance to cytotoxic therapies irrespective of niche, since these therapies target dividing cells. However, prognosis is likely shaped not by quiescence alone but by the surrounding ECM context. The limited remodeling in the Homeostatic ECM may maintain dormancy while the active remodeling Fibrous and Stress programs are more consistent with invasive niches and progression. QSCs may therefore be uniformly resistant to therapy but not uniformly prognostic, and their clinical impact may depend on the ECM program in which they are embedded.

Overall, this study is limited by the resolution of Visium spots, in which individual spots contain multiple cells and therefore cannot fully resolve cell-specific sources of ECM signals. However, because our analyses are focused on coordinated transcriptional programs rather than individual cell population, this limitation is unlikely to affect the identification of broad spatial ECM programs across specimens. Another limitation is the translation of spatial ECM programs to bulk transcriptomic datasets, which do not preserve spatial information. However, bulk transcriptomic profiling remains more applicable in clinical settings and therefore provides an important framework for evaluating the translational relevance of these findings.

In summary, this study identifies the ECM as a spatially organized and functional contributor to the GBM microenvironment. By defining distinct ECM transcriptional programs and linking them to different cell states, intercellular signaling and clinical outcomes, we demonstrate that ECM organization represents a vital axis of GBM biology. More specifically, the Homeostatic ECM, which has limited ECM remodeling, had the strongest and most consistent association with the QSC state, with the FGF1-FGFR axis emerging as a potential mechanism through which ECM programs may support QSC maintenance. Furthermore, elevated Homeostatic ECM expression in bulk transcriptomics was associated with improved survival following recurrence. Overall, these findings provide a framework for understanding how ECM-defined microenvironments contribute to GBM persistence and suggest how ECM-associated signaling pathways shape GBM biology, cellular plasticity, and clinical outcomes.

## METHODS

### Study Cohort

We analyzed a cohort of 26 IDH-wild type GBM Visium spatial transcriptomics specimens from 18 patients assembled by Greenwald et al. [7] which consisted of 13 sections generated at University Hospital Zurich, Massachusetts General Hospital, and Brigham and Women’s Hospital and 13 sections generated by Ravi et al. [26] that were jointly processed, quality-filtered, and annotated for CNA, Ivy GAP and genome-wide metaprograms. For the survival analysis we used the Glioma Longitudinal AnalySiS (GLASS) cohort.

### Consensus Non-Negative Matrix Factorization of ECM Gene Expression Programs

cNMF was applied to a curated ECM gene set from the matrisome database. Since factorization was restricted to an ECM gene set rather than genome-wide highly variable genes, and to ensure consistency with downstream analyses performed on normalized expression values, cNMF was conducted using log-normalized data, consistent with established use of log-normalized NMF applications for transcriptomic program discovery [27]. Prior to factorization, ECM genes were filtered to require detection in at least 20% of spots in at least 30% of specimens across the cohort resulting in 213 ECM genes across 60,189 spatial transcriptomics spots.

cNMF was performed using candidate factorization ranks of 2-6, with 200 iterations per rank and a fixed random seed for reproducibility. Consensus matrices were generated for each rank using local density filtering threshold of 0.01 to exclude poorly correlated replicate spectra, following the default cNMF outlier-filtering procedure. The optimal number of programs was selected using the cNMF k-selection diagnostic (**Fig. S1**), which balances reconstruction error against consensus stability. Consensus stability remained near maximal (>0.99) across k=2-4 before declining at k=5. Based on this stability plateau along with the interpretability and biological coherence of the resulting programs, k=3 was selected for downstream analyses.

cNMF decomposes the ECM expression matrix *V_g×n_* into a gene-weight matrix *W_g×k_* and a program usage matrix *H_k×n_* such that *V≍ WH*. Program usage values from the *H* matrix were used for analyses including spatial mapping, program spatial analysis, and associations with hallmark signatures. For survival analyses in bulk transcriptomic cohorts, ECM program scores were calculated using genes with weights greater than 1.5 standard deviations above the mean within each program in the *W* matrix.

### Spatial transcriptomics datasets and sample annotations

Spot-level malignancy annotations were obtained from copy number alteration (CNA), chromosome 7 gain and chromosome 10 loss. CNA-based annotations were assigned as defined by Greenwald et al. [7], classifying spots as *malignant*, *non-malignant*, *mix_high* or *mix_low*. Mixed-high (*mix_high*) spots were defined as spots with malignancy scores between the 50th and 75th percentiles of the malignancy-score distribution. For downstream analyses, neoplastic spots were defined as malignant and *mix_high*. Specimens containing at least 20% malignant or *mix_high* spots were considered in the malignancy restricted analyses, including NicheNet, and the neoplastic cell-cycle association analyses.

### ECM program assignment across specimens

Program usage scores *H* were mapped to the individual transcriptomics spots by matching spot identifiers. Program activities were incorporated into the metadata of each specimen for visualization and spatial statistical analysis. ECM program activity was visualized using spot-level coordinates extracted from each Visium specimen. To improve visualization and minimize the influence of extreme values, color scales were clipped at the 2^nd^ and 98^th^ percentiles of each program score distribution, with values outside this range squished to the nearest boundary.

### Differential gene expression analysis

Since Visium samples were not evenly distributed across patients and multiple samples were available for some patients, raw counts were aggregated at the patient level to avoid pseudoreplication. Counts were aggregated across all spots with the same dominant ECM program, generating one pseudo-bulk expression profile per patient-program combination, excluding any combinations with fewer than 20 spots. Differential gene expression analysis was performed with a single model fit across all three programs and accounting for inter-patient variability using the design *formula ∼ patient_id + program*. Genes were filtered to those with at least 10 counts in at least 25% of pseudo-bulk samples. Using that fitted model, three pairwise contrasts were extracted (Stress vs. Homeostatic, Homeostatic vs. Fibrous, Stress vs. Fibrous).

### Gene set enrichment analysis

For each pairwise contrast, genes were ranked by their differential gene expression Wald statistic and used as input for gene set enrichment analysis (GSEA) against the Hallmark, PID, and Reactome gene set collections in MSigDB. Significant pathways identified across pairwise comparisons were combined and their similarity was quantified using Jaccard distances based on overlap of GSEA core-enrichment genes. To reduce redundancy, representative pathways were selected from groups of highly similar pathways. Disease-specific or biologically unrelated pathways were removed from visualization to improve interpretability.

### Global spatial autocorrelation analysis

Spatial autocorrelation of ECM programs was quantified per specimen using Moran’s I statistic. Spatial neighborhoods were constructed using k-nearest-neighbor graph(k=6), corresponding to the hexagonal architecture of Visium. Row-standardized spatial weights, such that the weights assigned to all neighbors of a given spot summed to one, were used for all spatial autocorrelation analyses. For each ECM program, global Moran’s I was calculated and resulting p-values were adjusted across all tests using the Benjamini-Hochberg method. Positive Moran’s I values indicate spatial clustering, while values approaching 0 indicate spatial randomness.

### Spatial coherence analysis

Local spatial structure was examined using Local Moran’s I (LISA) statistics which breaks down global spatial autocorrelation into location-specific contributions. For each ECM program and specimen, Local Moran’s I values were computed and individual spots were assigned to one of four spatial association categories: High-High; elevated program activity surrounded by elevated neighboring activity, Low-Low (reduced program activity surrounded by reduced neighboring activity), High-Low: elevated activity surrounded by reduced neighboring activity. Low-High: reduced activity surrounded by elevated neighboring activity. Only spots with nominal significance (p <0.05) were assigned these classes, while all remaining spots were categorized as non-significant.

### Cross-program spatial association analysis

Spatial relationships between ECM programs were also assessed using Lee’s L spatial cross-correlation statistic. Pairwise analyses per specimen were performed for three ECM program combinations: Homeostatic ECM versus Stress ECM, Homeostatic ECM versus Fibrous ECM and Stress ECM versus Fibrous ECM. For each specimen, Lee’s L was calculated to quantify whether spatially neighboring regions associated with one program were also associated with another program. Positive values indicate spatial co-localization of paired ECM programs, while negative values indicate spatial segregation. P-values were adjusted using Benjamini-Hochberg method. To evaluate inter-specimen variability in ECM spatial architecture, specimens were clustered using Euclidean distance and complete-linkage hierarchical clustering based on their three pairwise Lee’s L statistics. Clustered heatmaps were generated to visualize similarities in ECM spatial organization across specimens.

### Non-ECM signatures quantification

All signatures were quantified per spots across specimens using Seurat’s AddModuleScore function. The Hippo pathway signatures were derived from a manually curated YAP/TAZ-Hippo transcriptional gene set. An immune suppression signature was defined from canonical immunosuppressive myeloid markers including MSR1, NT5E, VSIR, CD163, GPNMB, TREM2, and CSF1R. Focal adhesion genes were obtained from the KEGG Focal Adhesion pathway through MSigDB, while hypoxia genes were obtained from the MSigDB Hallmark Hypoxia gene set. Genes shared between the matrisome database genes and the query microenvironmental signatures were removed to prevent circular associations driven by shared genes. For all statistical modeling, genes included in the curated ECM gene set were removed from all non-ECM signatures prior to module score calculation to avoid circular associations between ECM programs and the target biological signatures.

### Univariate linear mixed models

For individual ECM program associations, separate univariate linear mixed-models were fitted for each ECM program-signature pair with random intercepts nested by patient and specimen to account for non-independence of spots within the same specimen and patient. Regression coefficients (β) and marginal R^2^ (R^2^m) values from these models were visualized as a dot plot, with color indicating the coefficients and point size indicating the proportion of variance explained.

### Nested mixed-effects models comparing ECM programs to histological annotation

To evaluate whether ECM programs explained biological variance beyond histological compartmentalization, nested mixed-effects models were applied to four non-ECM biological signatures (Hypoxia, Hippo, Immune Suppression, Adhesion) using all spots. For each spot, ECM program usage *H* were extracted for all three programs, and spatial metadata were integrated with Ivy GAP anatomical annotation. Both ECM program usage scores and functional signature scores were standardized with z-score transformation prior to modeling. For each signature, a full model containing Ivy GAP region and the three ECM programs was compared to a reduced model containing Ivy GAP region alone with specimen and patient identifiers nested as random intercept in both models to account for non-independence of spots within the same specimen and patient.

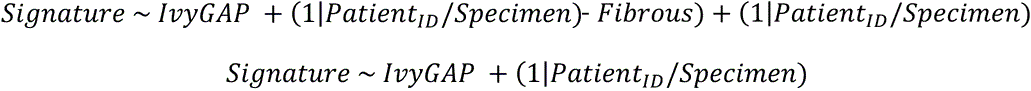

Models were compared via likelihood ratio test and Bayesian Information Criterion (BIC), which penalizes model complexity by the number of estimated parameters, with lower BIC values indicating better-supported models. The proportion of variance explained by fixed effects R^2^m was calculated using the method of Nakagawa and Schielzeth [28]. ΔR^2^m was defined as the increase in R^2^m when comparing the reduced (Ivy GAP-only) and full (Ivy GAP + ECM) models.

### Tertile visualization of ECM-signature associations by region

To visualize histological region-specific ECM-signature relationships, spots were divided into tertiles (Low, Medium, High) of each ECM program’s usage score, calculated separately within each Ivy GAP region. Mean signature score and standard error of the mean were calculated for each region × ECM tertile combination.

### Interaction likelihood ratio test across all spots

To test whether ECM program-signature associations vary by histological region, a linear mixed-effects interaction model (signature ∼ ECM × histologic, with random effects nested by patient and specimen) was compared against a main-effects-only model (signature ∼ ECM + histologic, with random effects nested by patient and specimen) via a likelihood ratio test. Region-specific slopes and 95% confidence intervals were extracted from the interaction model using estimated marginal trends (emtrends). A slope was considered significant if its 95% confidence interval did not include zero. *Neoplastic GBM states definitions*

QSC activity was quantified using the 118 gene set defined by Xie et al. [21]. For proliferation, a G2M-phase signature was used [29]. Glioma Stem Cell (GSC) activity was quantified using a curated GSC signature comprising stem and progenitor markers including core neurodevelopmental transcription factors reported to be functionally sufficient for GBM stem-like reprograming, together with additional established GSC markers spanning cytoskeletal protein linked to cellular stemness (NES) [30], an RNA-binding protein that helps control stem cells growth(MIS1) [31], epigenetic regulators promoting stemness (BM1,EZH2) [32, 33], stemness regulators (STAT3, FOXM1) [34, 35] and canonical stemness surface markers PROM1/CD133 [36]. Spatial transcriptomic spots were restricted to neoplastic regions. After filtering non-neoplastic spots and restricting the analyses to predominantly neoplastic specimens, a single specimen remained per patient, eliminating the need to model patient identifiers as nested random effects.

### ECM programs associations with cellular states in neoplastic spots

Analyses of tumor cell states were restricted to the neoplastic spots for specimens containing at least 20% neoplastic spots. ECM program usage scores were integrated with QSC, GSC and proliferation states after removal of ECM genes from each signature. Associations between ECM programs and cellular states were evaluated using the same univariate, nested mixed-effects, interaction, and tertile-based modeling framework described above, with specimen ID used as a random effect, after confirming that neoplastic specimens included only include one specimen per patient.

### ECM program-specific NicheNet analysis

NicheNet analyses were used to identify ECM-derived ligands predicted to regulate neoplastic QSCs across specimens. Analyses were performed independently for each of the three ECM programs. A QSC score was first calculated for neoplastic spots using the QSC signature after removing genes present in the ECM gene set to minimize circularity between ECM and quiescence measurements. Specimens were considered eligible if high-quiescence spots represented at least 10% of spatial spots, and at least 20 high- and 20 low-quiescence neoplastic spots were present following stratification.

NicheNet was performed separately for each eligible specimen. Sender spots were defined as spots with ECM program usage scores in the upper quartile ( ≥75^th^ percentile) of that program within the specimen. Receiver spots were restricted to neoplastic spots and stratified using a specimen-specific tertile split of the QSC score, while spots in the middle tertile were excluded from the receiver spots definition. For the quiescence group definition, we used a tertile stratification to have an adequate amount of receiver spots for optimal differential expression testing. ECM program-high sender spots were defined using a more stringent quartile threshold to prioritize specificity in identifying high-confidence signaling sources. To construct a consensus QSC-associated transcriptional program across specimens, differential expression analysis was performed independently within each eligible sample by comparing high and low QSC neoplastic spots using a Wilcoxon rank-sum test. Genes significantly upregulated (adjusted P < 0.05) in high QSC spots with detectable expression in at least 10% of the spots and a minimum log-fold change of 0.1 were saved per specimen. A consensus quiescence target gene set was then generated by selecting the genes that were identified in at least 50% of the eligible specimens. This consensus target set was used for all subsequent ligand activity analyses, ensuring that ligand activity predictions were evaluated against a common biological program across the cohort.

Candidate ligands were required to be present among marker genes defining the corresponding ECM program, expressed in ECM program-high sender spots, and connected through the NicheNet ligand-receptor network to at least one receptor expressed in high QSC receiver spots. Receptors were only considered if detected in 10% of receiver spots. Ligand activity scores were calculated using the NicheNet prior ligand-target model with the consensus QSC gene set as the target program and expressed receiver genes as the background set. Ligands were ranked within each specimen according to corrected area under the precision-recall curve (AUPR). To generate a consensus ranking across specimens, the mean corrected AUPR was calculated for each predicted ligand. The standard deviation of the corrected AUPR, mean rank and frequency of appearance among the top 10 ligands were also saved as measures of cross-specimen consistency. Ligands evaluated in less than 50% of specimens were excluded.

Spatial plausibility of predicted ECM-derived ligands was assessed to validate whether the ligands were expressed in the local microenvironment of QSC receiver spots. Mean ligand expression across the sender spots located within a defined neighborhood radius around each receiver spot was calculated. To avoid dependence on a single neighborhood radius definition, the analyses were repeated across a range of neighborhood radii corresponding to 2, 3, 4, 5, 6, 8, and 12 spot diameters. Cross-specimen consistency was summarized as the fraction of specimens exhibiting a positive localization across the tested neighborhood radii. Spatial stability was defined as positive localization in at least 70% of the neighborhood radii evaluated. A ligand was also required to show a net positive average difference in localized expression between high- and low-quiescence neighborhoods, pooled across specimens, to exclude genes whose apparent stability was driven by a narrow majority of specimens with opposing effect sizes (e.g. EDIL3). Final ligands were chosen based on both their mean AUPR rank in NicheNet activity analyses and their spatial stability fraction.

### Genome-wide metaprogram associations of predicted ligands

To investigate the cellular metaprograms associated with prioritized ligands while avoiding circularity, the metaprogram signatures defined by Greenwald et al. [7] were rescored after removing ECM-program associated genes. Each spot was then assigned to the metaprogram with the highest module score based on the resulting ECM-depleted signatures.

For each candidate ligand (FGF1, BGN), the analysis was restricted to spots belonging to the ligand’s producing ECM program, specifically, Homeostatic-high for FGF1 and Fibrous-high for BGN. For each specimen *s* for each metaprogram *m*, a metaprogram expression contrast was calculated as

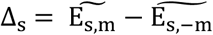

Where 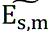 is the mean ligand expression among spots assigned to metaprogram *m* and 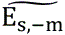 is the mean expression among all remaining metaprograms within the same ECM expressing spots. To assess consistency across specimens, a one-sample Wilcoxon signed-rank test was applied to the per-specimen Δ values to test whether the median contrast differed from zero across specimens. Resulting p-values were adjusted for multiple testing using the Benjamini-Hochberg method. Metaprograms with significantly positive Δ values were interpreted as cellular states in which ligand expression was consistently associated with the metaprogram across specimens.

### ECM programs scoring in GLASS cohort

RNA sequencing data and clinical annotations from GLASS were obtained through Synapse. Transcript-level count data were mapped to gene symbols and collapsed to gene-level expression by summing counts across transcripts corresponding to the same gene. For samples with multiple sequencing portions from the same specimen, raw counts were averaged to generate a single expression profile per case and timepoint. Library-size normalization was then performed using counts per million (CPM) followed by log-transformation. All survival analyses were restricted to IDH-wild type GBM patients with primary and first recurrence RNA transcripts available for longitudinal analyses.

ECM program activity was quantified using single-sample gene set enrichment analysis (ssGSEA). Genes exceeding the mean by 1.5 standard deviations in the Homeostatic, Stress and Fibrous ECM programs were used to define program-specific gene sets. ssGSEA enrichment scores were calculated independently for each sample using normalized gene expression profiles. Changes in ECM program activity between matched primary and recurrent tumors were evaluated using paired Wilcoxon signed-rank tests and resulting p-values were adjusted for multiple testing using the Benjamini-Hochberg method.

### Longitudinal change in ECM transcriptomic levels

To evaluate longitudinal transcriptomic changes between the primary tumor and the recurrent tumor, ssGSEA scores were compared between matched pairs using Wilcoxon signed-rank tests. Median changes in program activity were calculated for each ECM program, and p-values were adjusted for multiple testing using the BH approach.

### Bulk transcriptomics survival analysis

Two clinical endpoints were evaluated: (1) time from primary surgical resection to first recurrence and (2) survival following recurrence, defined as the interval from recurrence surgery to death or last follow-up (overall survival – time to recurrence). For each clinical endpoint, three Cox models were compared for discriminative performance: a clinical model with age, sex, and MGMT promoter methylation status as covariates, and ECM-only model, and a combined model containing both clinical covariates and the ECM program scores. Performance across these models was assessed using 5-fold cross-validation with a ridge penalizer of λ= 0.1, and a mean out-of-fold concordance index (C-index) was reported for each model.

To estimate the independent prognostic value of ECM programs after adjustment for clinical covariates, unpenalized multivariate Cox regression models (clinical-only and combined) were fitted to the full cohort with hazard ratios (HRs), 95% confidence intervals, and Wald test p-values reported.

### Statistical packages

All analyses were conducted in R and Python. Spatial transcriptomic processing and visualization were performed using Seurat, RcppML, presto, NMF, dplyr, ggplot2, viridis, patchwork, cowplot, Matrix, and related packages. Ligand-receptor inference was performed using the nichenetr package. Survival analyses, including Cox proportional hazard regression, Kaplan-Meier estimation, and log-rank testing, were implemented using the Python package lifelines. ECM program activity in bulk transcriptomic data was quantified using gseapy. Additional analyses used pandas, NumPy, SciPy, statsmodels, atplotlik, and scikit-learn. Multiple-testing adjustment was performed using the Benjamini-Hochberg false discovery rate method. Adjusted two-sided p-values < 0.05 were considered statistically significant.

## Supporting information

Supplemental Figures used for the link to the file on the preprint site

## Acknowledgements

This work was supported by the US National Institutes of Health National Cancer Institute P50CA221747, U54CA302435, R01CA295560, R01CA247905, National Institute of Neurological Disorders and Stroke U24NS133949, National Library of Medicine award R01LM013523, and DOD Glioblastoma Resource Development Grant 14301880

## Author Contributions

M.A.A, D.J.B conceived the research idea. M.A.A carried out data analysis, model development, and wrote the paper. L.A.D.C provided expertise in bioinformatics and machine learning. L.S, P. B, and H.N provided expertise in spatial transcriptomics. J.L. provided expertise on neurooncology. M.A.A, L.A.D.C, P.B. H.N, and D.J.B edited the paper. D.J.B supervised the work.

