## Supplemental Figures used for the link to the file on the preprint site for "Extracellular matrix spatial ecosystems define stem-like niches and predict post-recurrence survival in glioblastomas"

### ABSTRACT

Extracellular matrix (ECM) remodeling is a hallmark of glioblastoma (GBM) yet has typically been studied through a reductionist lens, as individual genes or pathways rather than large scale, spatially integrated analyses coordinated with biological states. Focusing on whole matrisome gene expression, we identified three reproducible and spatially coherent ECM transcriptional programs, termed Homeostatic, Stress, and Fibrous, that were associated with specific microenvironmental functions and neoplastic cell states, with the strongest association noted between Homeostatic ECM and neoplastic quiescent stem cell-like (QSC) niches. Ligand-receptor analyses identified FGF1 and BGN as candidate ECM-driven regulators of QSC. In an independent longitudinal bulk transcriptomic cohort, ECM-based models outperformed traditional clinical and biomarker variables for predicting post-recurrence survival, with elevated Homeostatic ECM activity associated with improved post-recurrence survival, while MGMT promoter methylation no longer remained prognostic. Overall, these findings establish ECM organization as a clinically relevant dimension of GBM biology and nominate ECM-associated signaling as candidate therapeutic targets.

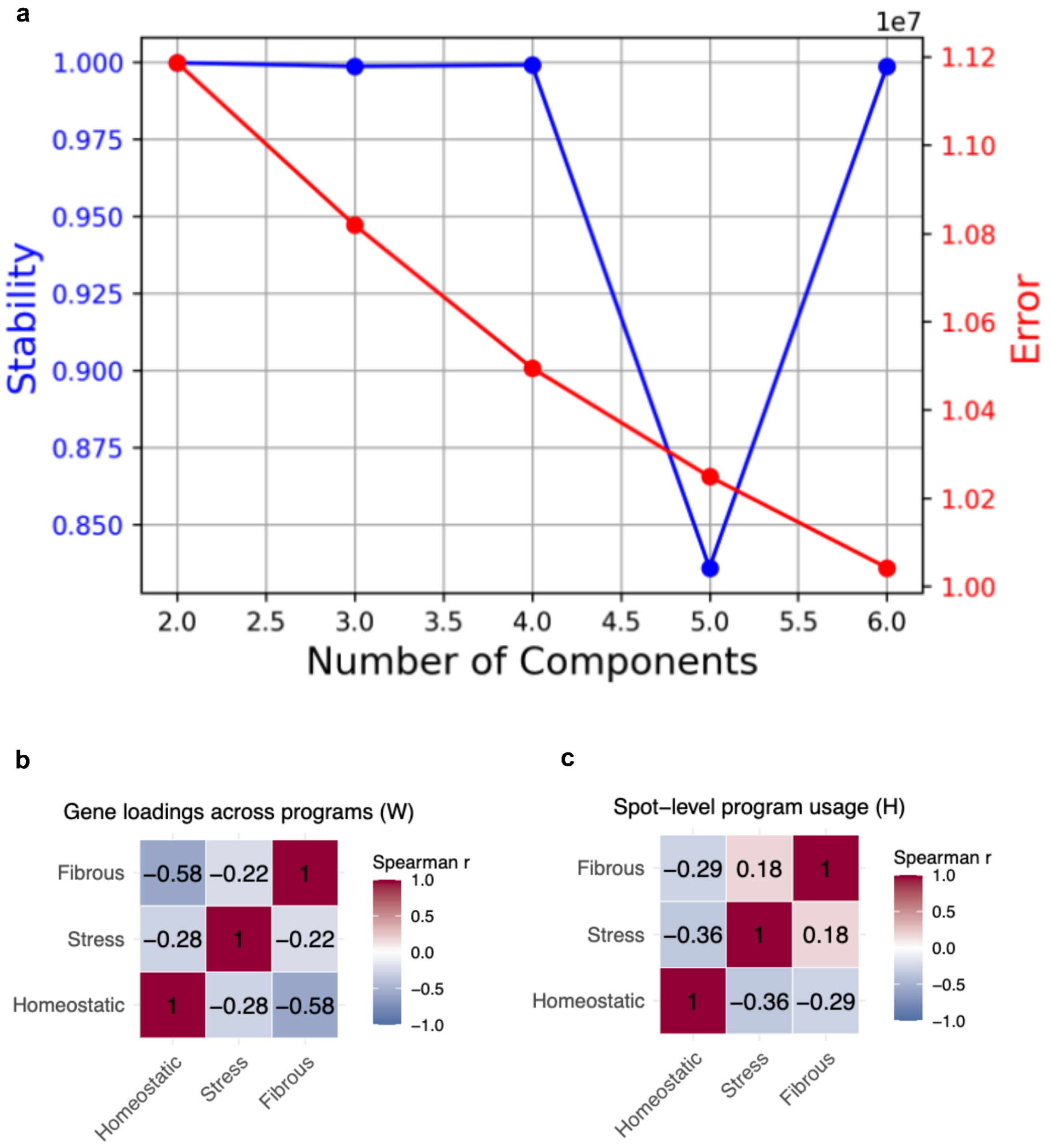


**Figure S1. Selection and validation of cNMF-derived transcriptional programs. (a)** k-selection diagnostics for cNMF (applied to matrisome gene expression across Visium spots. Reconstruction error (red) and consensus stability (blue) were evaluated across factorization ranks (k = 2-6), with k=3 selected balancing model fit and stability. **(b)** Spearman correlation of gene loadings (W) between the three ECM programs, demonstrating distinct program-specific genes. **(c)** Spearman correlation of spot-level ECM program usage scores (H), showing independent patterns of program activity across spatial transcriptomic spots.

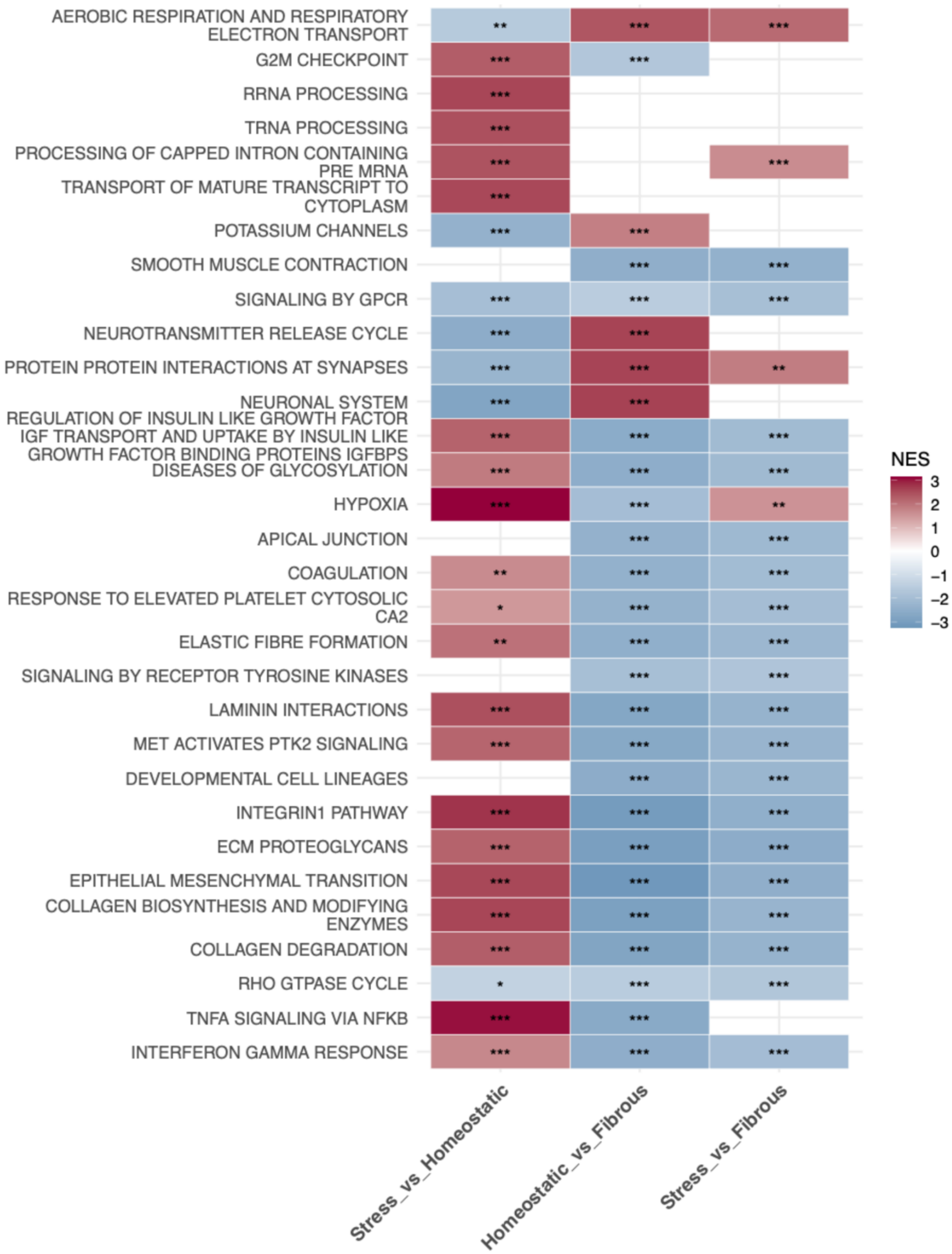


**Figure S2. Pairwise GSEA**. Complete set of molecular pathways enriched across ECM programs. Color scale represents normalized enrichment score (NES). Color scale represents normalized enrichment score (NES); asterisks denote adjusted p-value (BH-corrected)(*p < 0.05, **p < 0.01, *** p < 0.001, blank tiles not significant)

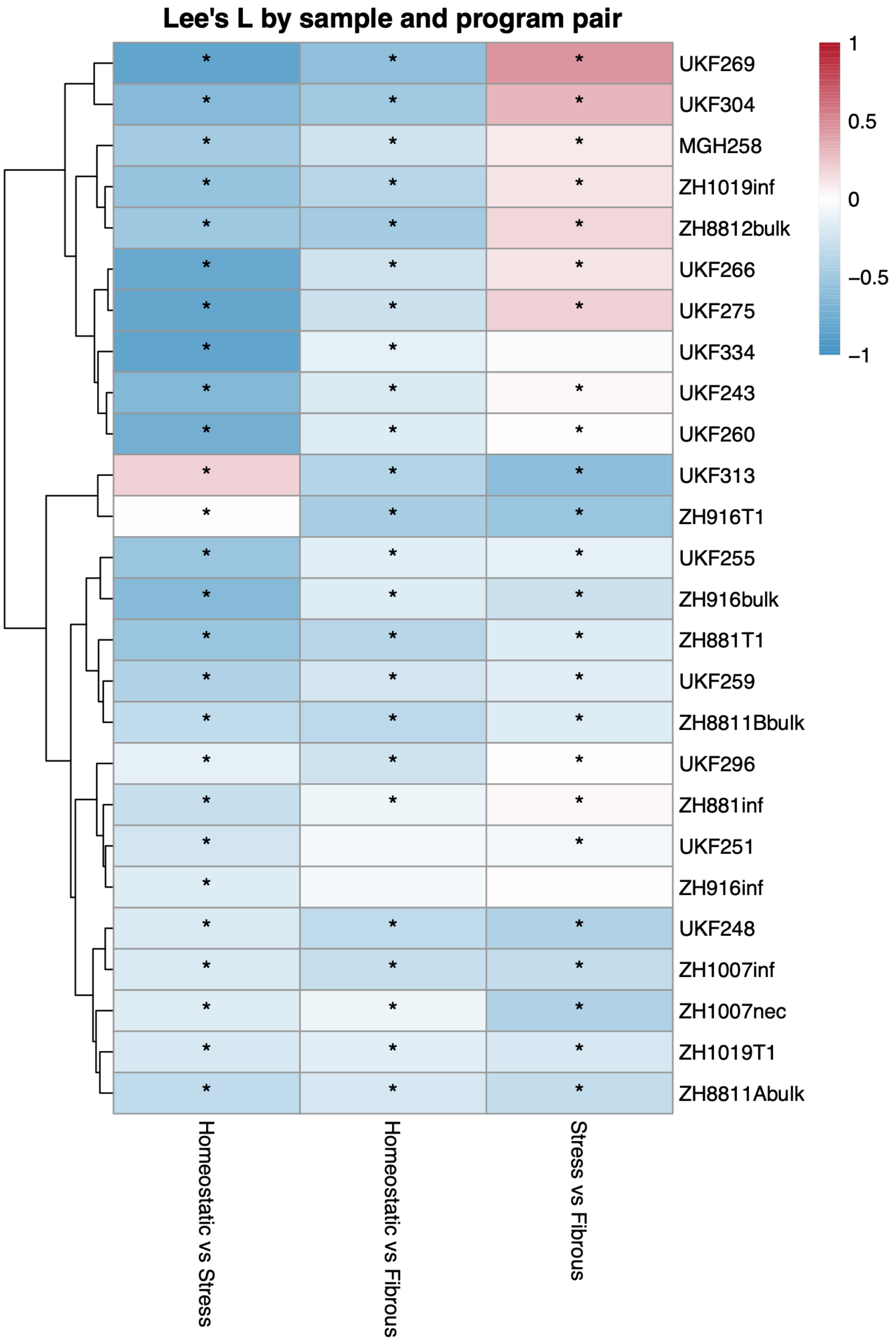


**Figure S3. Spatial associations between ECM transcriptional programs across specimens**. Heatmap showing Lee's L spatial statistic for pairwise ECM program combinations across specimens. Positive Lee's L values (Red) represent ECM program pair colocalization. Negative Lee's L values (Blue) represent ECM program pair segregation. Asterisks denote statistically significant spatial associations ($p < 0.05$). Specimens were hierarchally clustered based on Lee's L values. Homeostatic is mostly segregated from Fibrous and Stress ECM, Stress and Fibrous showed positive spatial associated in the top cluster.

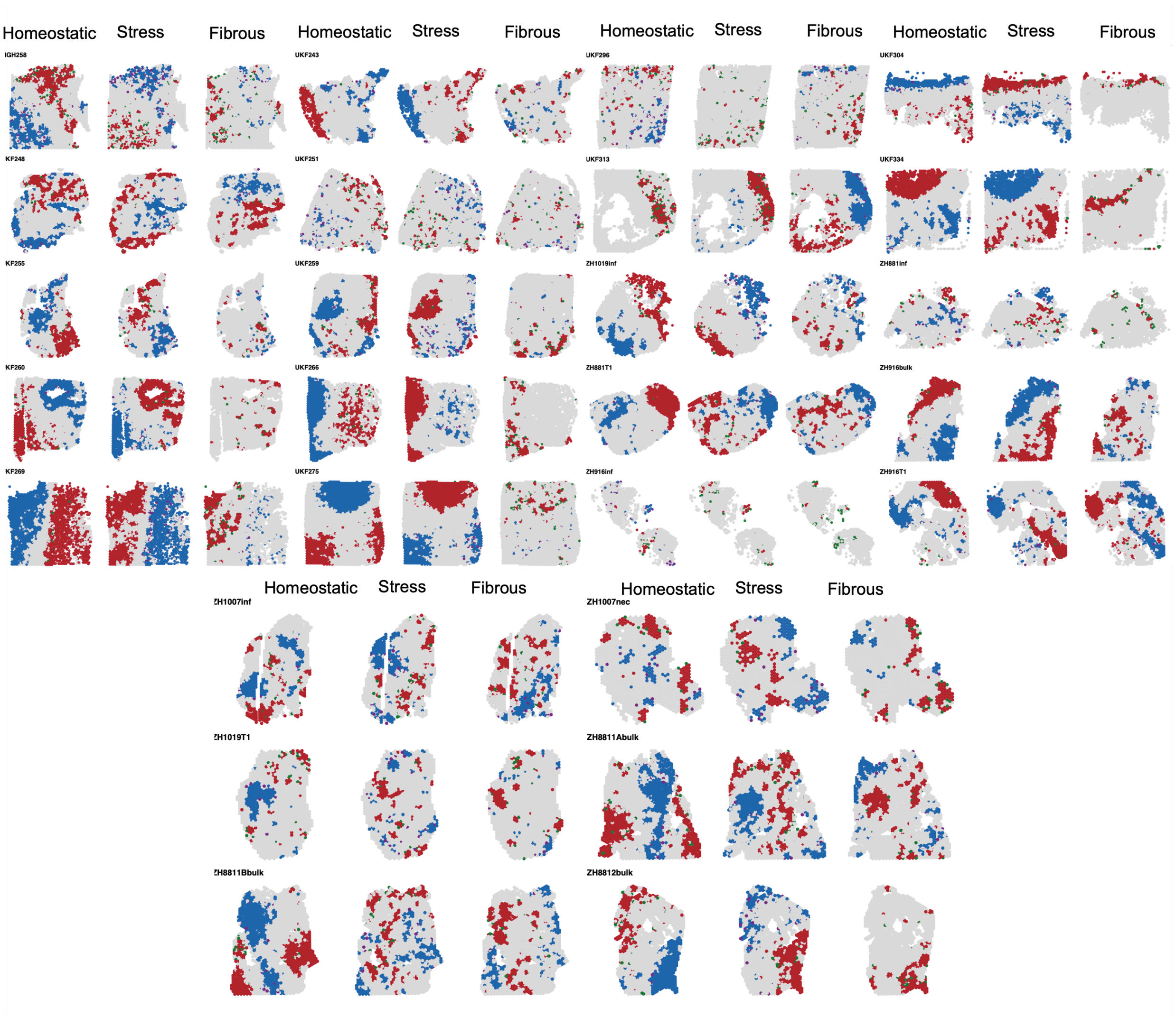


**Figure S4. LISA maps for cohort.** Local spatial association (LISA) maps showing regions of ECM hot spots (red), cold spots (blue), transitioning neighborhoods: from high to low(purple), from low to high(green), non-significant (NS) spatial autocorrelation (gray)

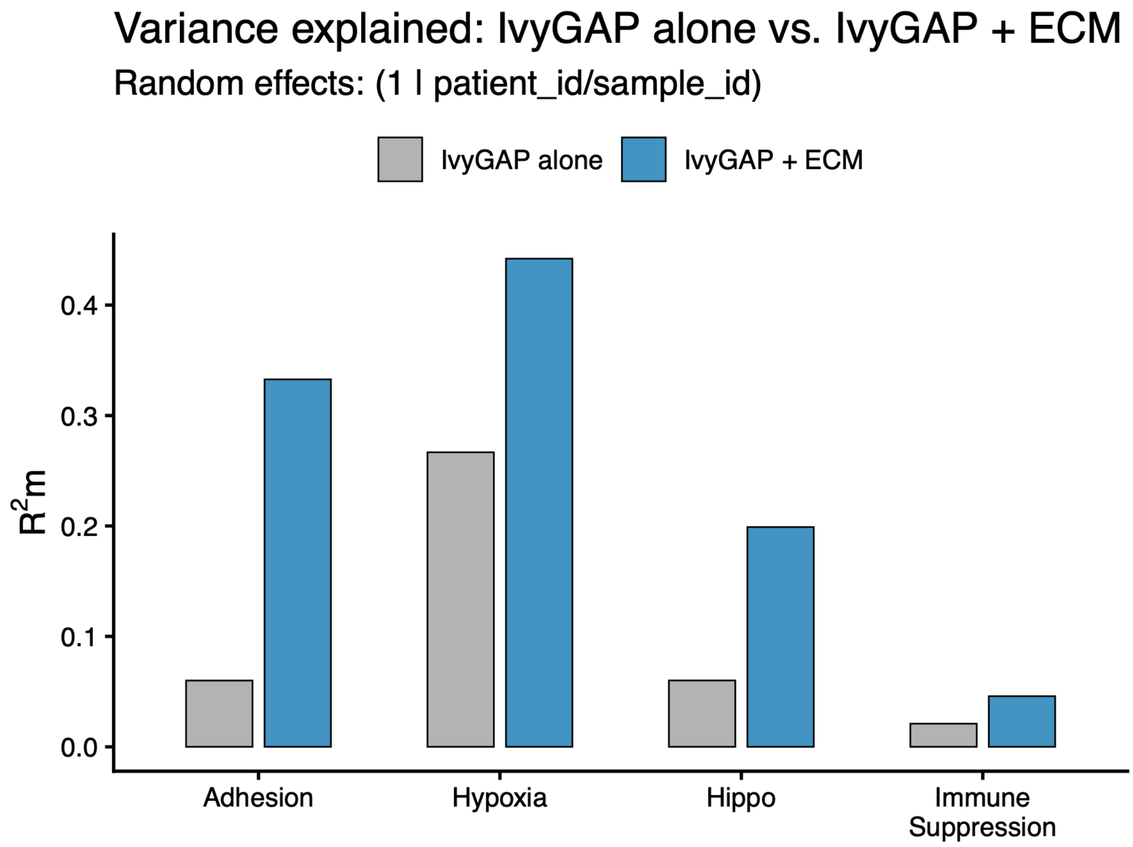


**Figure S5. Comparison of variance explained between histological classification and ECM programs**. Marginal $R^2m$ from linear mixed-effects model predicting hallmark signatures using Ivy GAP histological regions alone (gray) and a combined model with ECM program scores (blue).

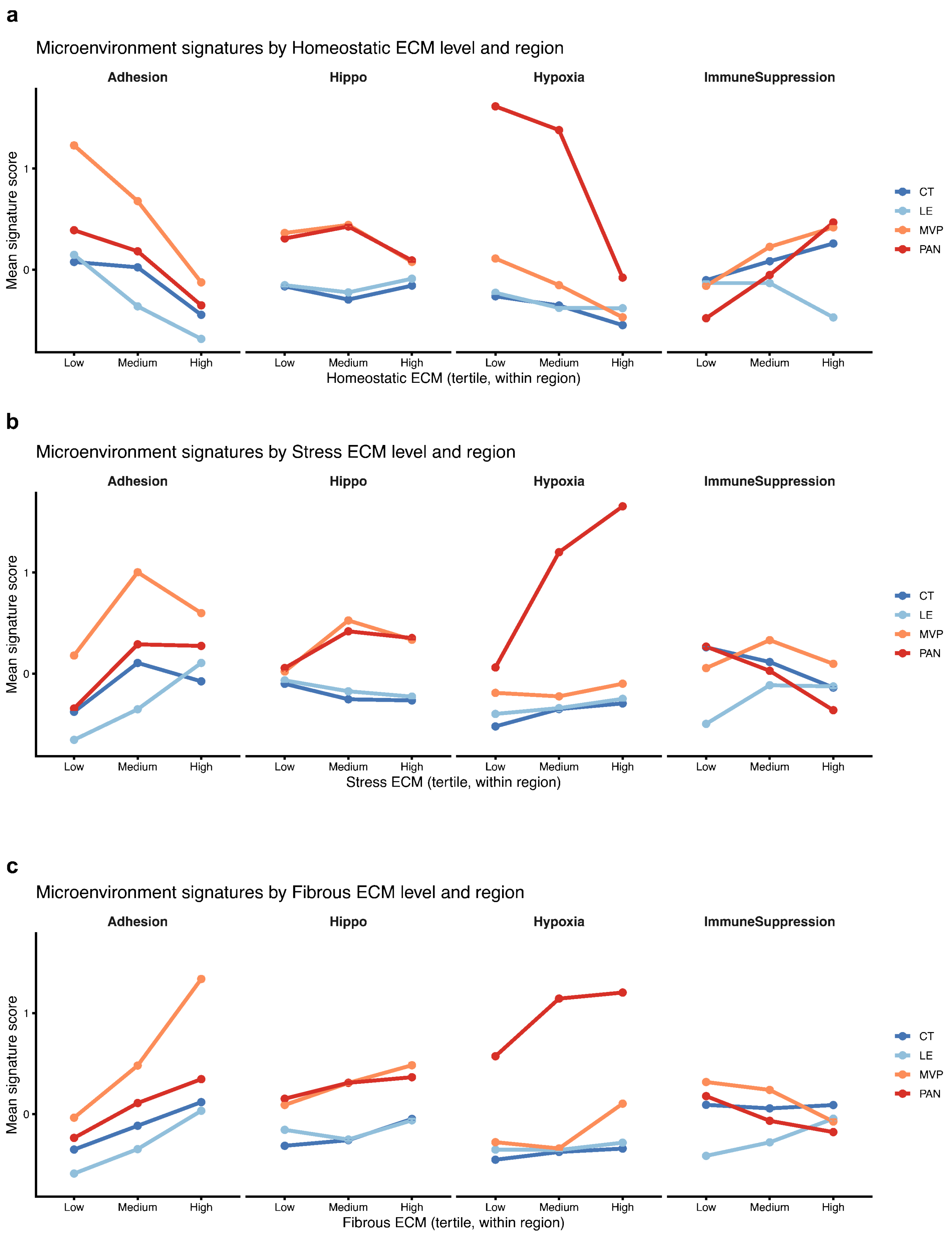


**Figure S6. Tertile ECM activity trends across histological regions**. Mean microenvironment signature scores across Ivy GAP histological regions stratified by ECM program activity tertiles **(a)** Homeostatic, **(b)** Stress, **(c)** Fibrous ECM programs

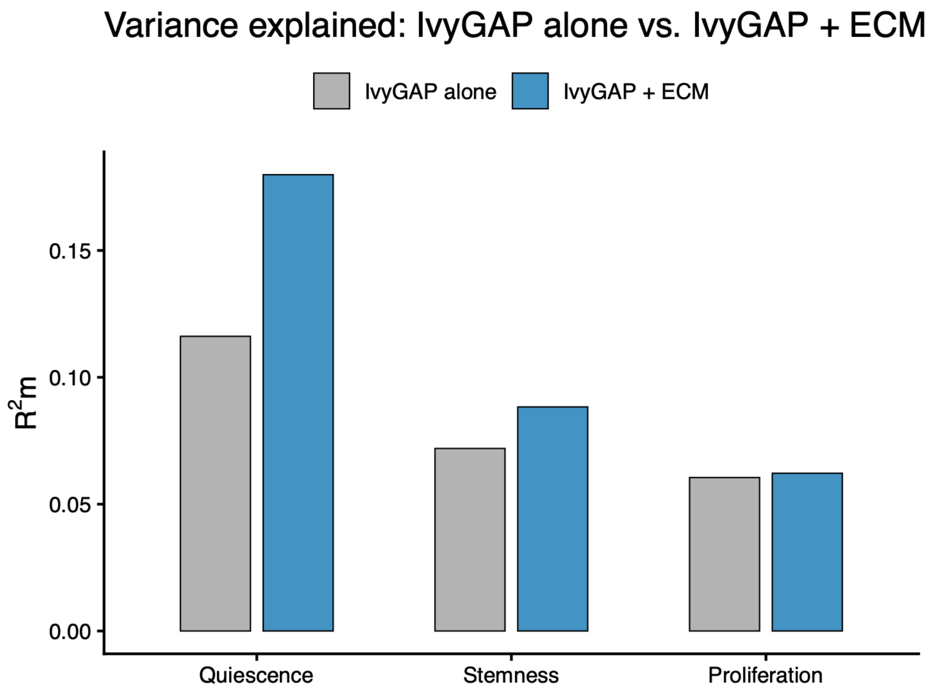


**Figure S7. Comparison of variance explained between histological classification and ECM programs for neoplastic cell states**. Marginal $R^2$m from linear mixed-effects models QSC, GSC and G2M proliferation signatures using Ivy GAP histological regions alone (gray) and a combined model incorporating Ivy GAP regions and ECM program scores (blue). Inclusion of ECM programs increased the variance explained for QSC and GSC, with only a minimal increase for G2m proliferation.

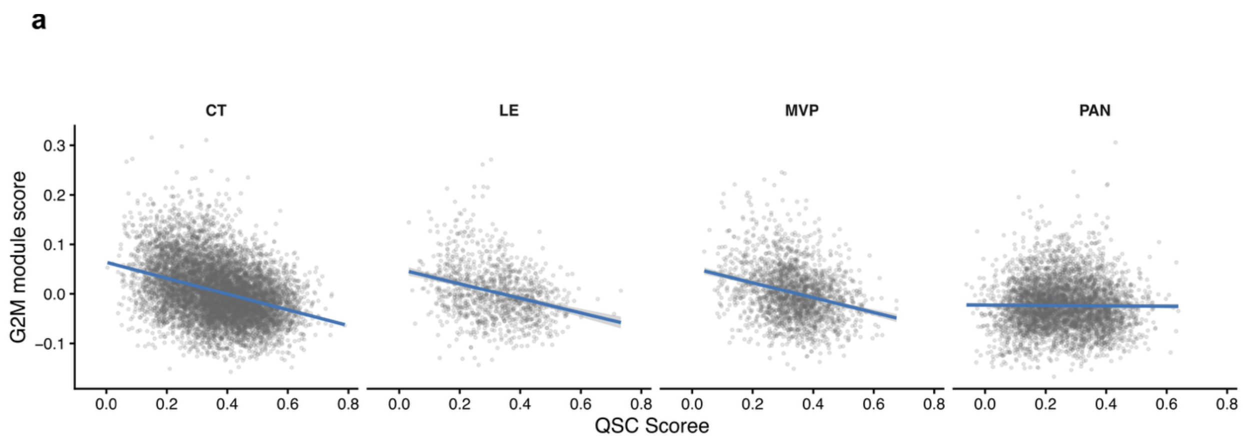


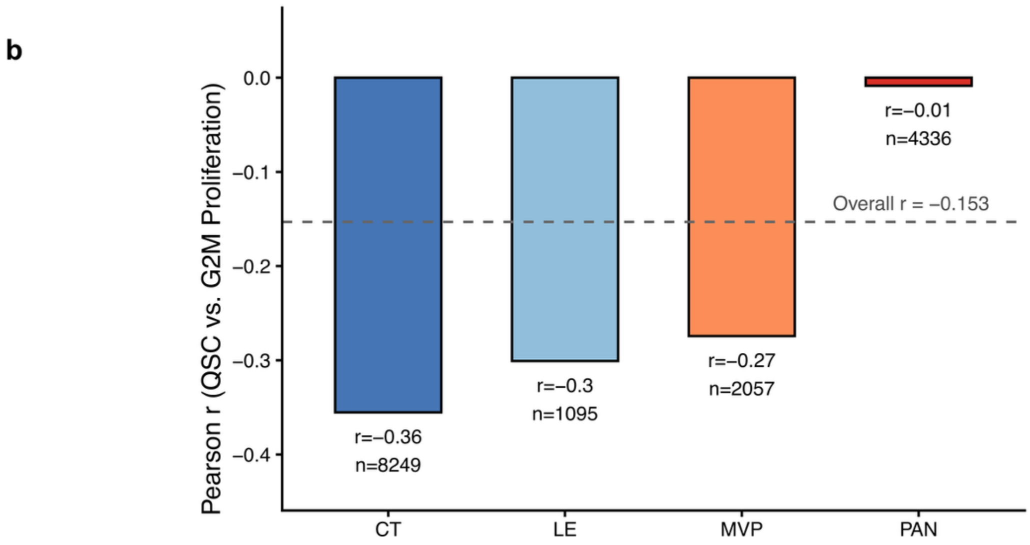


**Figure S8. QSC and G2M proliferation anticorrelation across histological regions**. (a) Scatterplots showing the relationship between QSC vs G2M proliferation scores stratified by region limited to neoplastic Visium spots. Blue lines are linear regression fits, showing negative correlation between QSC and proliferation across histological regions except for in the pseuadopalisading and perinecrotic regions (PAN). (b) Pearson correlation coefficients between QSC and proliferation scores stratified by region. *n* is the number of spots in each comparison. The dashed line denotes the overall correlation. Pearson's r shows minimal associations between QSC and proliferation in PAN.

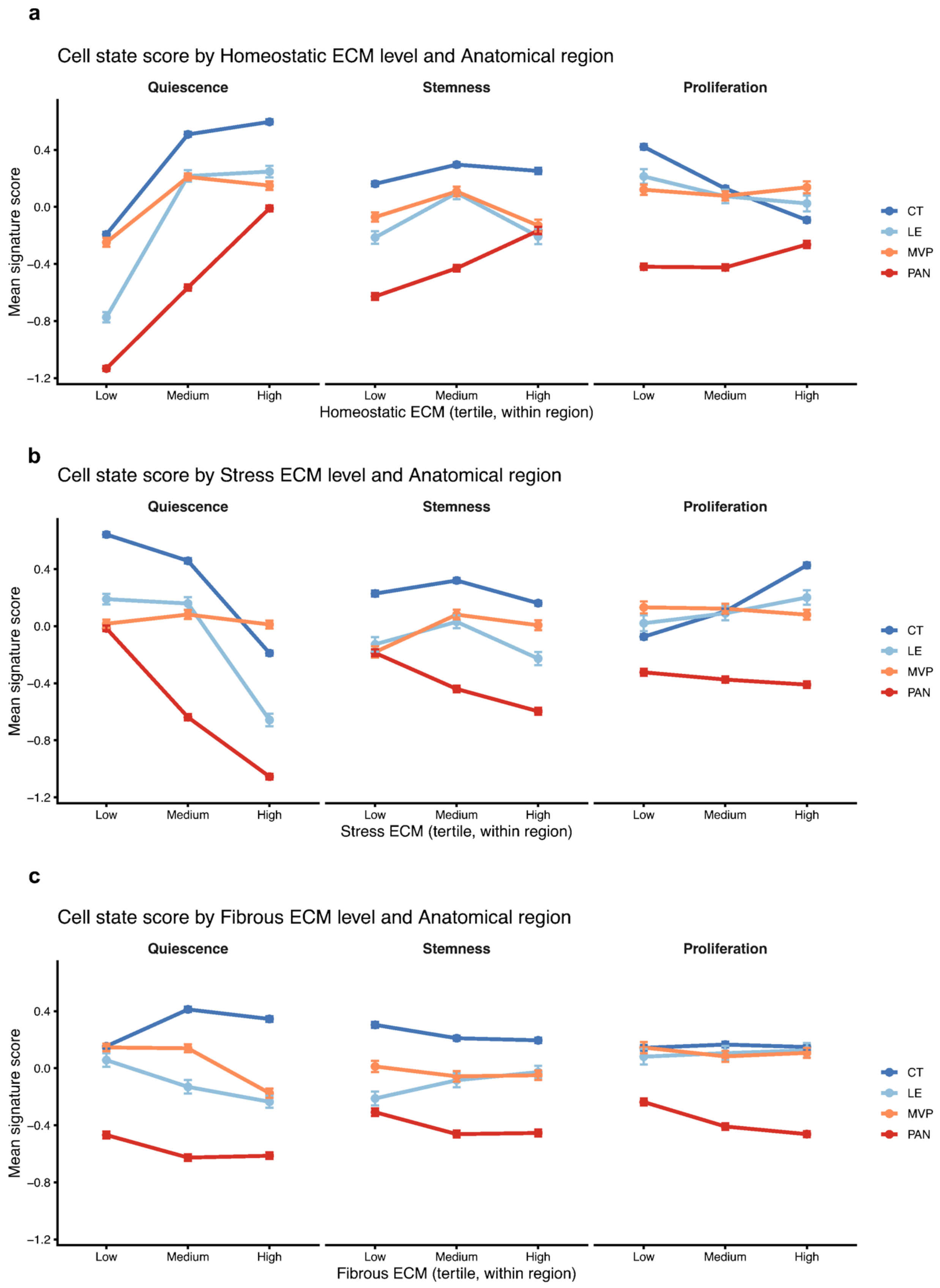


**Figure S9. Tertile ECM activity trends across histological regions for neoplastic spots**. Mean neoplastic cell state scores across Ivy GAP histological regions stratified by ECM program activity tertiles **(a)** Homeostatic, **(b)** Stress, **(c)** Fibrous ECM programs

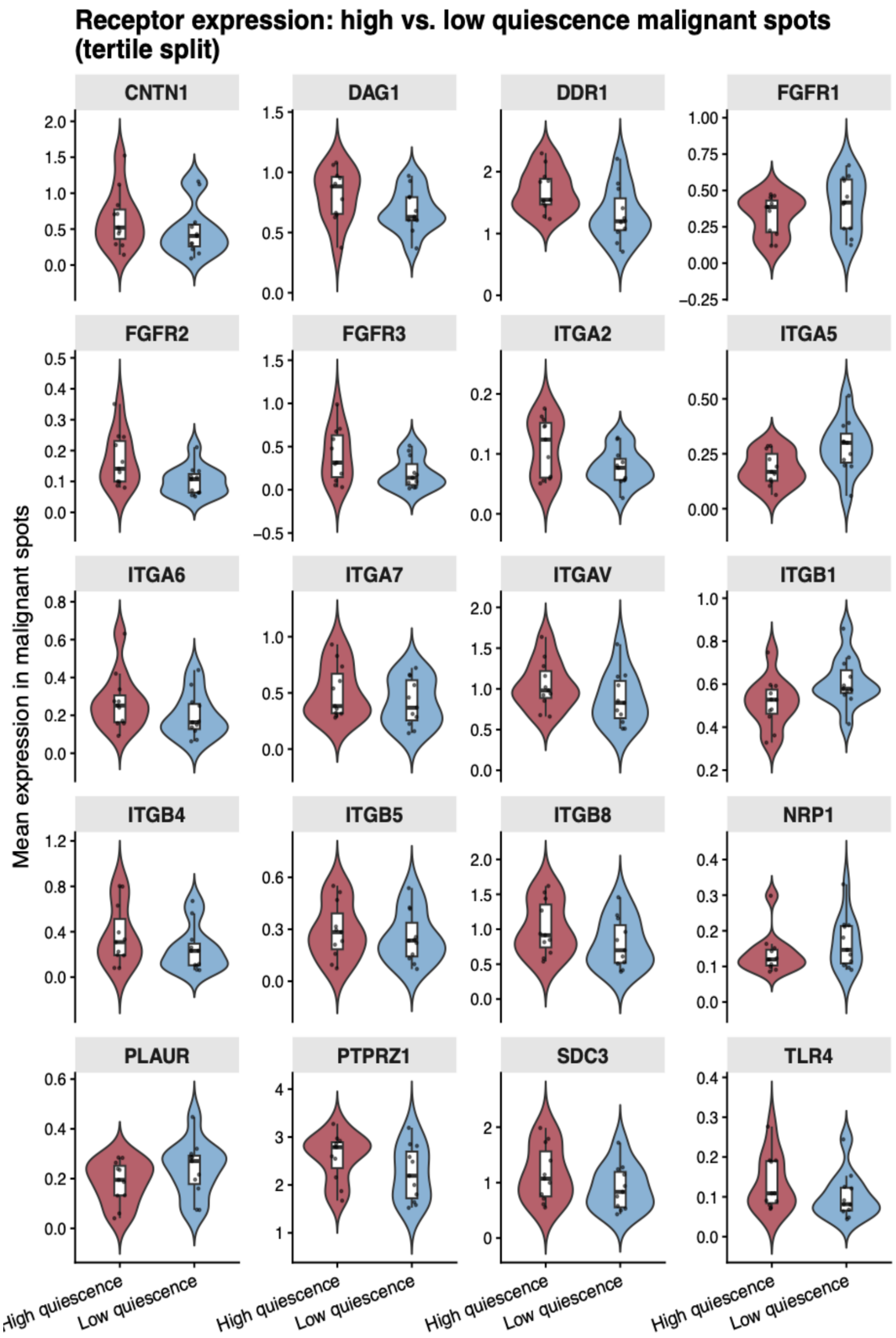


**Figure S10. Expression of predicted receptors in high- and low- quiescence neoplastic spots**. Expression of candidate receptors associated with ECM-derived ligand signaling in neoplastic spots stratified by QSC tertiles. For each specimen, mean receptor expression was calculated in high0 and low0- QSC spots and compared using paired Wilcoxon signed-rank tests across specimens. Only receptors significant after BH correction are show ($p < 0.05$).

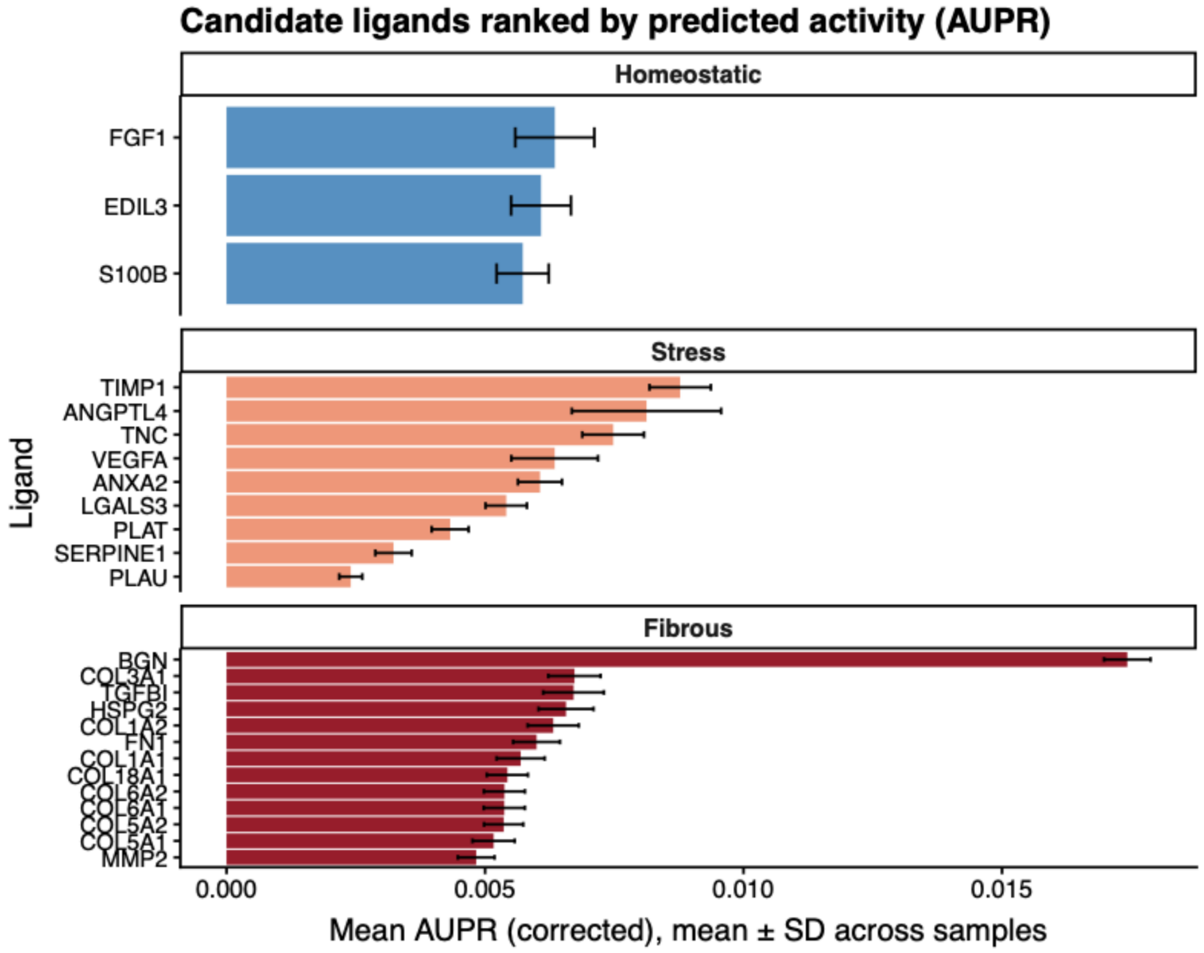


**Figure S11. Candidate ECM-derived ligands ranked by predicted activity**. NicheNet ligand activity scores for ECM programs using ECM-high spots as senders and QSC high neoplastic spots as receivers. Ligands are ranked by mean corrected AUPR across specimens with error bard indicating standard deviation. Higher AUPR values indicate stronger predicted regulation of QSC spots.

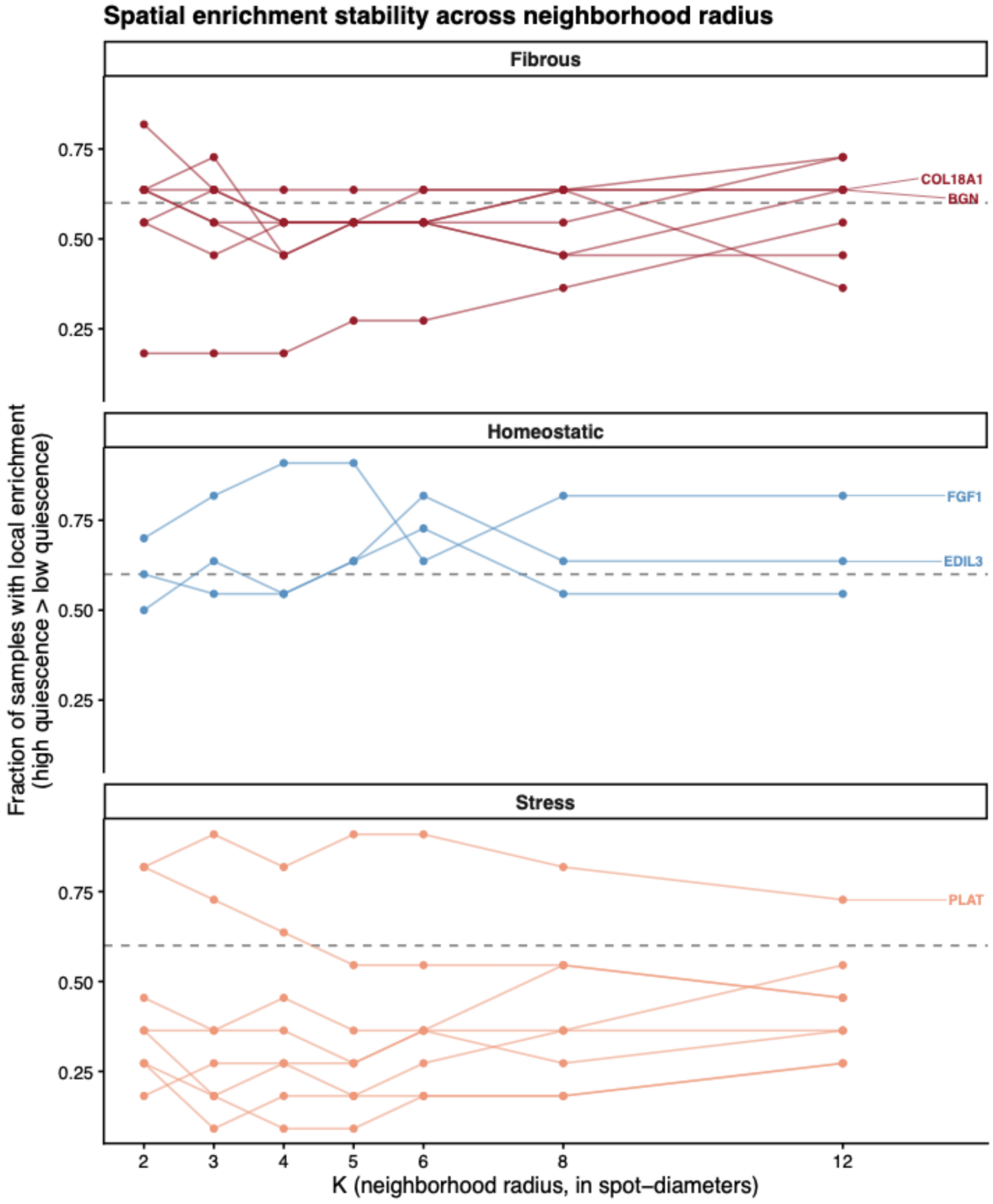


**Figure S12. Spatial stability of candidate ECM-derived ligands across neighborhood radii**. A sweep across neighborhood radii ranging from 2 to 12 spot diameters to examine spatial stability of candidate ligands identified by NichNet. For each ligand and radius, the fraction of specimens in which local ligand expression was greater around high than low QSC neoplastic spots. The dashed line indicated the enrichment threshold (60% of the specimens). Ligands consistently exceeding this threshold across multiple neighbor radii were considered spatially stable candidates.

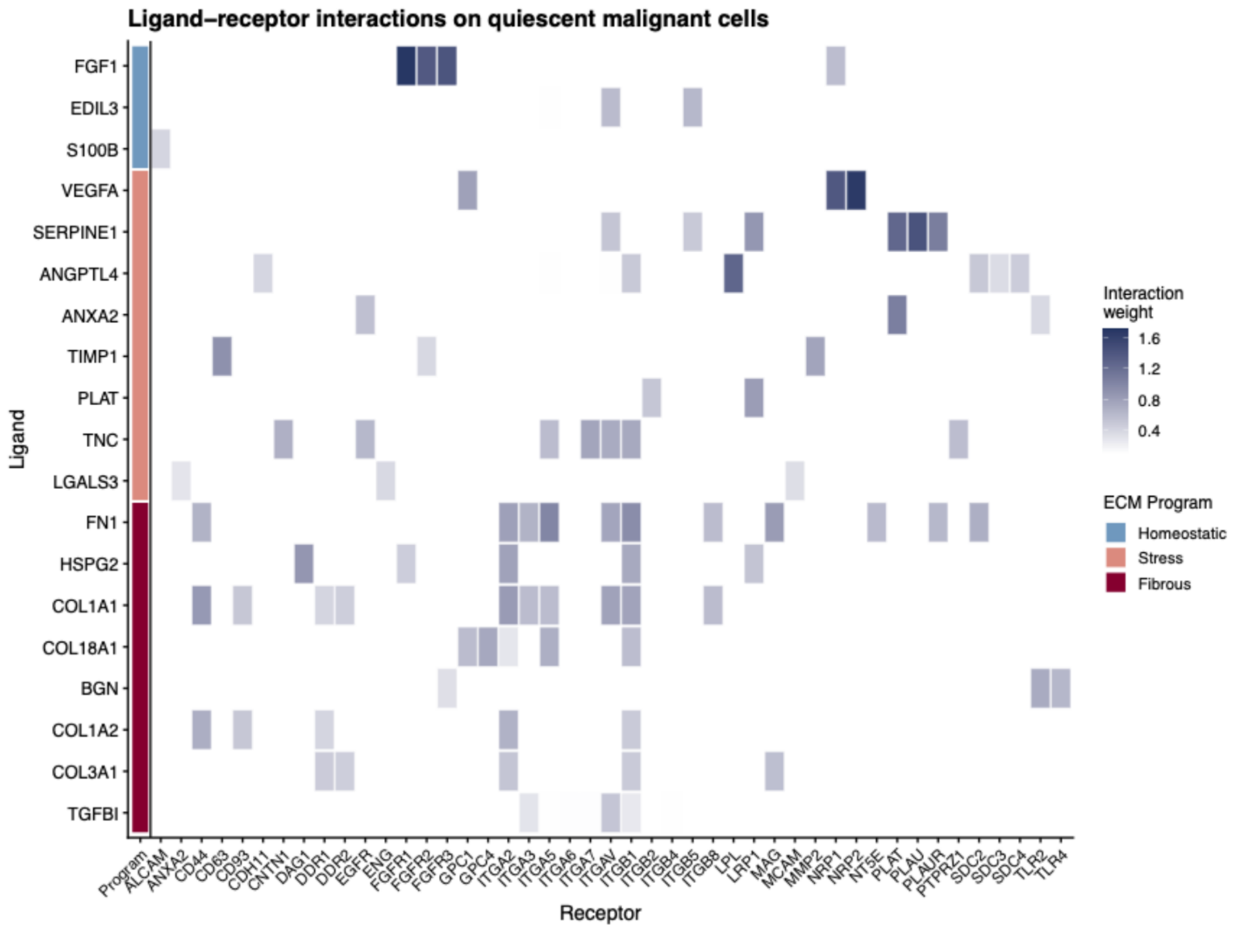


**Figure S13. Ligand receptor interactions on QSC neoplastic spots.** Heatmap showing predicted interactions between top-ranked NicheNet ECM program-ligands and receptors expressed in high QSC neoplastic spots. Color intensity represend ligand-receptor interaction weights derived from the NicheNet prior regulatory network.

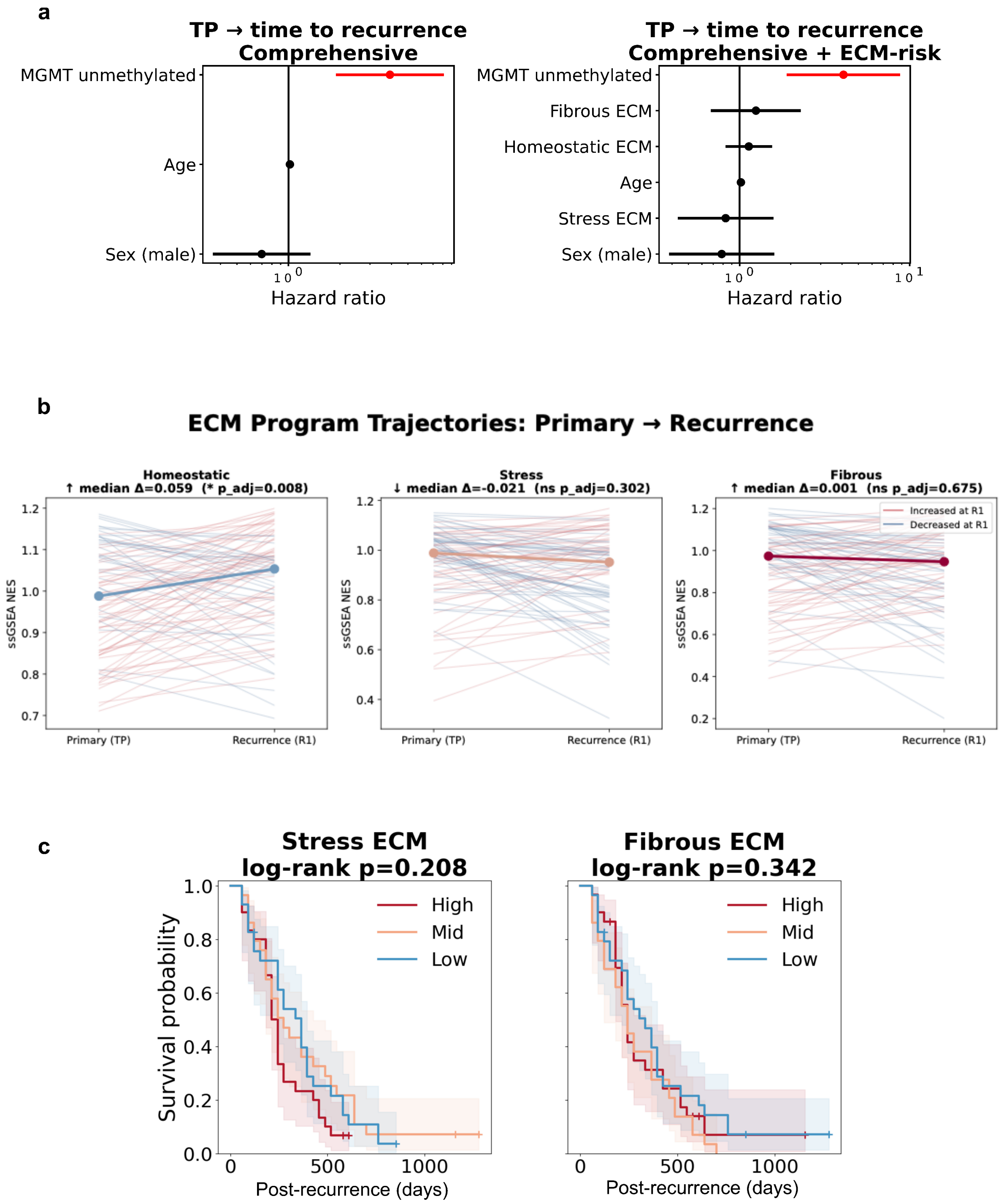


**Figure S14. ECM program dynamics and survival associations in the GLASS cohort. (a)** Multivariable Cox proportional hazards model for time to recurrence comparing clinical only with a combined model adding ECM programs activities. Hazard ratios and 95% confidence intervals are shown with red line indicating significance ($p < 0.05$). MGMT promoter methylation status remained the strongest prognostic variable in recurrence prediction. **(b)** Longitudinal changes in ECM program activity between matched primary and recurrent GBM specimens. P-values were calculated using Wilcoxon signed-rank tests. Homeostatic ECM activity showed increase at recurrence, while Stress and Fibrous ECM activity remained unchanged. **(c)** Kaplan-Meier plots of post-recurrence survival stratified by tertiles of recurrent Stress and Fibrous ECM programs. Shaded areas indicate 95% confidence intervals. Log-rank p-values show significant difference between groups ($p > 0.05$).
